# Machine learning-guided directed evolution for the development of small-molecule antibiotics originating from antimicrobial peptides

**DOI:** 10.1101/2022.11.03.515123

**Authors:** Heqian Zhang, Yihan Wang, Pengtao Huang, Yanran Zhu, Xiaojie Li, Zhaoying Chen, Yu Liu, Jiakun Jiang, Yuan Gao, Jiaquan Huang, Zhiwei Qin

## Abstract

Antimicrobial peptides (AMPs) are valuable alternatives to traditional antibiotics that possess a variety of potent biological activities by exerting immunomodulatory effects to clear difficult-to-treat infections. Understanding the structure-activity relationships (SARs) of AMPs can direct the synthesis of desirable therapeutics. In this study, we use machine learning-guided directed evolution to develop the lipopolysaccharide-binding domain (LBD), which acts as a functional domain of anti-lipopolysaccharide factor (ALF), a family of AMPs, identified from *Marsupenaeus japonicus*. We report the identification of LBD_A-D_ as an output of this algorithm with the input of the original LBD_*Mj*_ sequence and show the NMR solution structure of LBD_B_, which possesses a circular extended structure with a disulfide crosslink in each terminus and two 3_10_-helices and exhibits a broad antimicrobial spectrum. Scanning electron microscopy and transmission electron microscopy showed LBD_B_ induced the formation of a cluster of bacteria wrapped in a flexible coating that ruptured and consequently killed the bacteria. The co-injection of LBD_B_ and *Vibrio alginolyticus, Staphylococcus aureus* and another major pathogen in shrimp aquaculture white spot syndrome virus *in vivo* improved the survival of *M. japonicus*, indicating a promising therapeutic role of LBD_B_ for infectious disease. The findings of this study pave the way for the rational drug design of activity-enhanced peptide antibiotics.

## Introduction

Antimicrobial resistance (AMR) caused by the misuse of traditional antibiotics is one of the greatest medical challenges and most alarming issues for public health in the 21^st^ century(*1*). AMR leads to approximately 700,000 deaths each year, and this number could increase to 10 million by 2050 if new antibiotics or therapies fail to develop(*2*). Despite the urgent need for new antibiotics with new targets, for commercial and technical reasons, there are only a few candidates currently in clinical trials. More than half of currently used antibiotics are derived from microbial natural products and were introduced into clinics during the golden age between 1940 and 1960. Since then, discoveries have declined as the results of repeated isolation of the already known compounds. Although large-scale genome sequencing in recent years has shown that microbial sources are still a highly promising reservoir for the discovery of novel bioactive natural products as antibiotic leads, the bottleneck of isolation mentioned above can arguably be attributed to the fact that the cultivation of most microorganisms (> 99%) and natural product biosynthesis (> 80%) are hampered under laboratory conditions(*3, 4*). Thus, many efforts have been made to provide new approaches for the expression of biosynthetic gene clusters for natural product discovery(*5*). However, this process cannot compete effectively against the progression of AMR. Compared to these natural product-oriented antibiotics, which are biosynthesized via secondary metabolism and typically called ‘small molecules’ due to their relatively low molecular weights, antimicrobial peptides (AMPs) represent a promising alternative large class of therapeutic agents with higher molecular weights (generally between 20 and 60 amino acids) and are derived from a variety of organisms, such as bacteria, plants, invertebrates and vertebrates, in response to the innate immune system(*6–8*). AMP exhibit broad-spectrum antimicrobial, antifungal and antiviral activities as well as immune regulation functions(*9–11*) and its activity usually associate with the presence of cationic and hydrophobic residues, such as arginine (R), lysine (K) or histidine (H) for the former and tryptophan (W), phenylalanine (F) or leucine (L) for the latter(*12*). These two types of residues have physicochemical features and biological functions that allow AMPs to mediate bacterial lipid interactions and cell membrane associations, eventually causing damage or disruption of the cytoplasmic membrane of microorganisms) (*13*). The mode of action of AMPs has been well proposed in recent years, and important mechanisms consist of pore formation(*14*), carpet-like(*15*) and barrel models(*16*). Other mechanisms include targeting cellular and metabolic processes, such as the biosynthesis of cell wall(*17*), protein(*18*), DNA(*19*) and RNA(*20, 21*) or the bioprocesses of enzymatic functions(*17*) and cell division(*22*).

Anti-lipopolysaccharide factor (ALF) is an essential innate immune defensive protein and a key immune effector in crustaceans(*23–26*). Several studies have shown that different isoforms of ALF coexist in one organism(*27, 28*). ALFs can be divided into seven groups (A to G) in penaeid shrimp according to their primary structures and biochemical characteristics(*29, 30*). The lipopolysaccharide-binding domain (LBD) is the key functional domain of ALF and usually consists of α-helix or β-hairpin structures, as well as a disulfide loop formed by two cysteine residues that aid the LBD in exerting the antimicrobial property of ALF(*31, 32*). Recent studies have shown that structurally modified LBDs can improve bioactivities and thus provide a promising reservoir for new antimicrobial agent design or optimization^(*33*),(*34*),(*35*)^. We have recently reported the ALF family AMP (ALF_*Mj*_) isolated from the kuruma prawn *Marsupenaeus japonicus*, which shows activity against *Photobacterium damselae* and *Micrococcus luteus*(*36*) however, its corresponding LBD_*Mj*_ did not generally exhibit activities when the experiments were performed in parallel, leaving a chance for further structural modification.

Given the advantages of a quick killing effect and high barriers to drug resistance, AMPs have attracted a lot of attention and interest in recent years. However, many AMPs are cytotoxic to mammalian cells, which hampers further development in clinical trials. A possible solution is structural modification in terms of the replacement of amino acids based on natural peptide sequences. Considering the nearly unlimited chemical space of AMPs (e.g., in theory even a 20-residue peptide can combine 20^20^ sequences), the widely available amino acids in nature, and the advantages of artificial peptide synthesis, it is necessary to develop a fast, cost-effective and accurate computing algorithm to accelerate the development of target AMPs with enhanced activity and less cytotoxicity. In recent years, scientists have developed specific algorithms for AMP bioengineering and have shown excellent applications. Yoshida *et al* reported a general linear model combined with a genetic algorithm and successfully identified 44 peptide sequences with markedly enhanced activity(*37*). Porto *et al* recently showed a genetic algorithm in which the α-helix index for fitness value was used and resulted in the target AMP guavanin 2, which kills the reporter strain *Escherichia coli* at lower concentrations)(*38*). Other illustrative examples include density-based clustering algorithms to identify AMPs active against gram-negative bacteria(*39*) and QSAR-based artificial neural network models for the identification of potent AMPs active against multi-drug resistant *Pseudomonas aeruginosa*(*40*). However, the common disadvantage of these algorithms appears to be large data volume or a high cost of iterative verification.

In this study, we develop a machine learning-based algorithm suitable for the directed evolution of LBD_*Mj*_. We first used fuzzy c-means clustering (FCM) to classify the published LBD sequences (10) performing excellent anti-*E. coli* activities with regard to net charge, hydrophobicity and isoelectric point, and found that they had markedly clustered populations on a three-dimensional plane. To explore the amino acid space for LBD_*Mj*_ substitution, we applied partial least squares regression (PLSR) on all currently published LBD sequences (30) and their MICs to construct a fitness matrix for roulette wheel selection. In addition, Monte Carlo simulation was conducted on the distribution of positively charged amino acids to derive more space for residues of LBD_*Mj*_ to be replaced with lysine, allowing us to establish an initial fitness matrix (equation) based on the collected LBD sequences mentioned above to generate directed optimized LBD sequences. Also, we combined relative-MIC value assignment and a genetic algorithm (GA) to iteratively enlarge the population of low relative-MIC values. Finally, the strong substitution sites were identified by the assembled machine learning algorithm proposed in this study and successfully validated the *in silico* optimized LBD sequences that can be used as guidance for chemical synthesis, NMR solution structure determination and systematic activity testing. In this work, we show that LBD_A-D_, the selected four sequences from the computing output, markedly enhance antimicrobial activities against *S. aureus, V. alginolyticus* and WSSV infection compared to the ‘wild-type’ LBD_*Mj*_. In addition, LBD_B_ shows good *in vivo* antiviral activity against WSSV, improving the survival of *M. japonicus.* Moreover, to evaluate the output from this algorithm, we randomly designed, synthesized and tested two peptides, LBD_X_ and LBD_Y_, that have identical amino acid lengths to LBD_A-D_ but different residues. Results show that the antimicrobial activities of LBD_X_ and LBD_Y_ were in between those of LBD_*Mj*_ and LBD_A-D_, indicating that the cationic and hydrophobic residues indeed play important roles in the activity of AMPs, but their numbers and positions in the peptide sequences must be considered carefully during optimization. To our knowledge, there are currently no such algorithms and applications for SAR enhancement for ALF family AMPs; therefore, the results of this study provide creative instructions for the design of LBD-originating antimicrobial agents and are valid for any other types of AMP modification.

## Results

### LBD sequence analysis

In this study, LBD_*Mj*_ produced from kuruma prawn *M. japonicus* was chosen as a study model. A BLAST search using this sequence identified another 30 sequences secreted by crustaceans, denoted LBD_1-30_ in this study (Supplementary Table 1). Among these sequences, the most similar one was LBD_9_ (truncated from FcALF-LBD6 produced by the Chinese shrimp *Fenneropenaeus chinensis*(*35*)) with 72.73% identity, while some others, such as LBD_8_ (LvALF8-LBD from the whiteleg shrimp *Litopenaeus vannamei)* or LBD_30_ (EcLBD4 from the Ridgetail Prawn *Exopalaemon carinicauda*), exhibit no marked similarities with LBD_*Mj*_. All sequences have a length of 22 amino acids and two cysteines in each terminus with a disulfide bridge. LBD_1-10_ exhibit potent activities against *E. coli*, and all have a MIC below 30 μM, which we consider to be potential for antibiotic drug development, while LBD_11-30_ was used as a negative reference because they have higher MICs. The alignment of LBD_*Mj*_ against LBD_1-10_ revealed the highly conserved Y^3^, V^5^, P^7^, L^14^, Y^15^, F^16^, and G^18^, while only R^11^ appeared to be conserved when LBD_*Mj*_ was blasted against LBD_11-30_. The average net charge, hydrophobicity, hydrophobic moment, and isoelectric point from LBD_1-10_ are 4.60 (±2.84), 0.53 (±0.08), 0.13 (±2.29), and 10.30 (±0.07), respectively, while those of LBD_11-30_ are 4.2 (±1.32), 0.57 (±0.14), 0.15 (±0.05), and 10.91 (±0.46), respectively. These results suggest that there are no apparent physicochemical differences between the two peptide groups, particularly two of the most important activity determinants-net charge and hydrophobicity, indicating that the simple sequence alignment could not allow accurate MIC prediction.

### Machine learning-guided directed evolution to optimize the structure of LBD for enhanced activity

In this study, we build a machine learning-based algorithm to assist in evolution-guided optimization rather than altering the net charge of an original peptide sequence, as shown in previous studies(*41, 42*). Thus, we used the above LBD_1-30_ sequences as the original dataset for algorithm development (Supplementary Table 1). As shown in Fig. 1, this algorithm consists of the construction and optimization module (COM) and strong substitution identification (SSI). The COM is described as below:

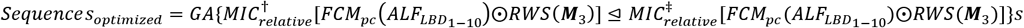

where 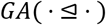 is the iterative optimization process guided by genetic algorithm, for example, 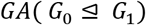 represents the direction of evolution from *G*_1_, to 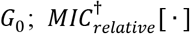 is the *in silico* LBD population with relative-MIC ≤ 15; 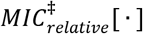 is the *in silico* LBD population with relative-MIC ≥ 15; *FCM_pc_*(·) is the fuzzy c-means clustering process based on physicochemical properties; *RWS*(***M***_3_) is the roulette wheel selection process based on the final fitness matrix ***M***_3_; ⊙ represent the relative-MIC value assignment and filtering process on *in silico* LBD population which is generated from *RWS*(***M***_3_).

**Fig. 1.**
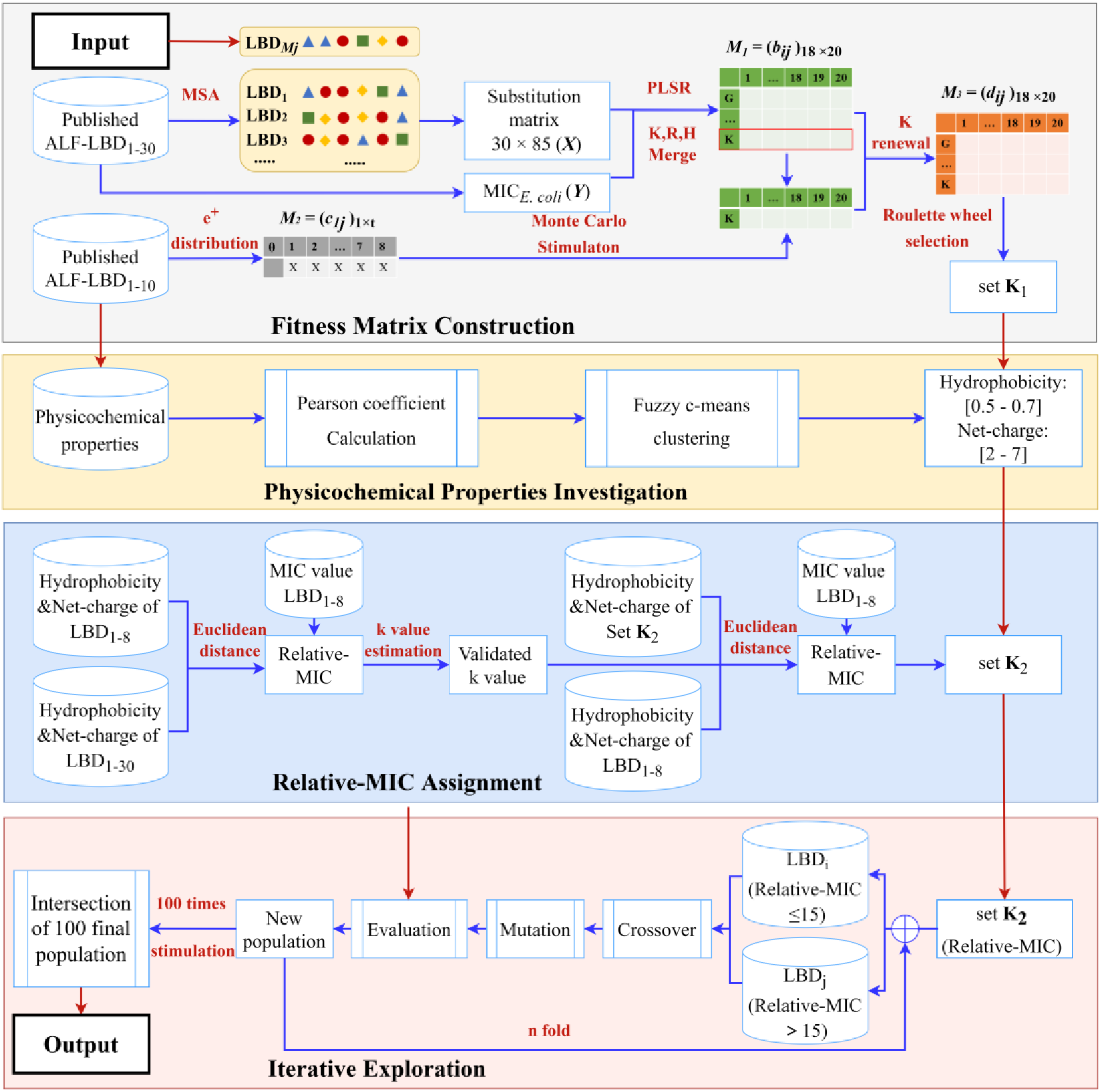
Illustration of the machine-learning-based algorithm in this study. The first part in this algorithm is construction and optimization module (COM) which accepts the input of original peptide sequences used in this study and comprises four subprocesses, including physicochemical property investigation, fitness matrix construction, relative-MIC assignment and iterative exploration, respectively. The second part is strong substitution identification (SSI) that has been embedded in COM, and this is the module for the output of optimized peptide sequences.

COM comprises four subprocesses, including physicochemical property investigation, fitness matrix construction, relative-MIC assignment and iterative exploration, respectively. In the process of physicochemical property investigation, the Pearson correlation coefficient measurement between MIC and physicochemical properties of the peptide dataset indicated that it is not practical to predict the MIC of a modified peptide sequence if simply comparing the difference between physicochemical properties of the original and modified peptides (Fig. 2a). Interestingly, three clusters were found using FCM with hydrophobicity, net charge and isoelectric point. LBD_1-8_ were the largest cluster created in space of hydrophobicity ranging from 0.5 to 0.7, net charge of 2 to 7, and isoelectric point of 10 to 12 (Fig. 2b and c). In the fitness matrix construction process, PLSR was first used to project the 30×85 substitution matrix and the corresponding MIC values to construct the initial fitness matrix ***M***_1_ (Fig. 2d). The final fitness matrix ***M***_3_ was then generated after the frequency distribution matrix ***M***_2_ of positively charged amino acids in each LBD_1-10_ was introduced to the fitness K, R, and H with Monte Carlo simulation (Fig. 2d). Thus, the matrix ***M***_3_, which described the PLS coefficient, was introduced to determine the probability of approximate 9432 newly generated LBD sequences (Set **K**_1_) in which three amino acids were randomly substituted (Supplementary Table 2). Then, set **K**_1_ was filtered by retaining only the preset net charge and hydrophobicity as indicated above, which resulted in a new library of optimized LBD sequences (Set **K**_2_) with a reduced number to 1593 (Supplementary Table 2).

**Fig. 2.**
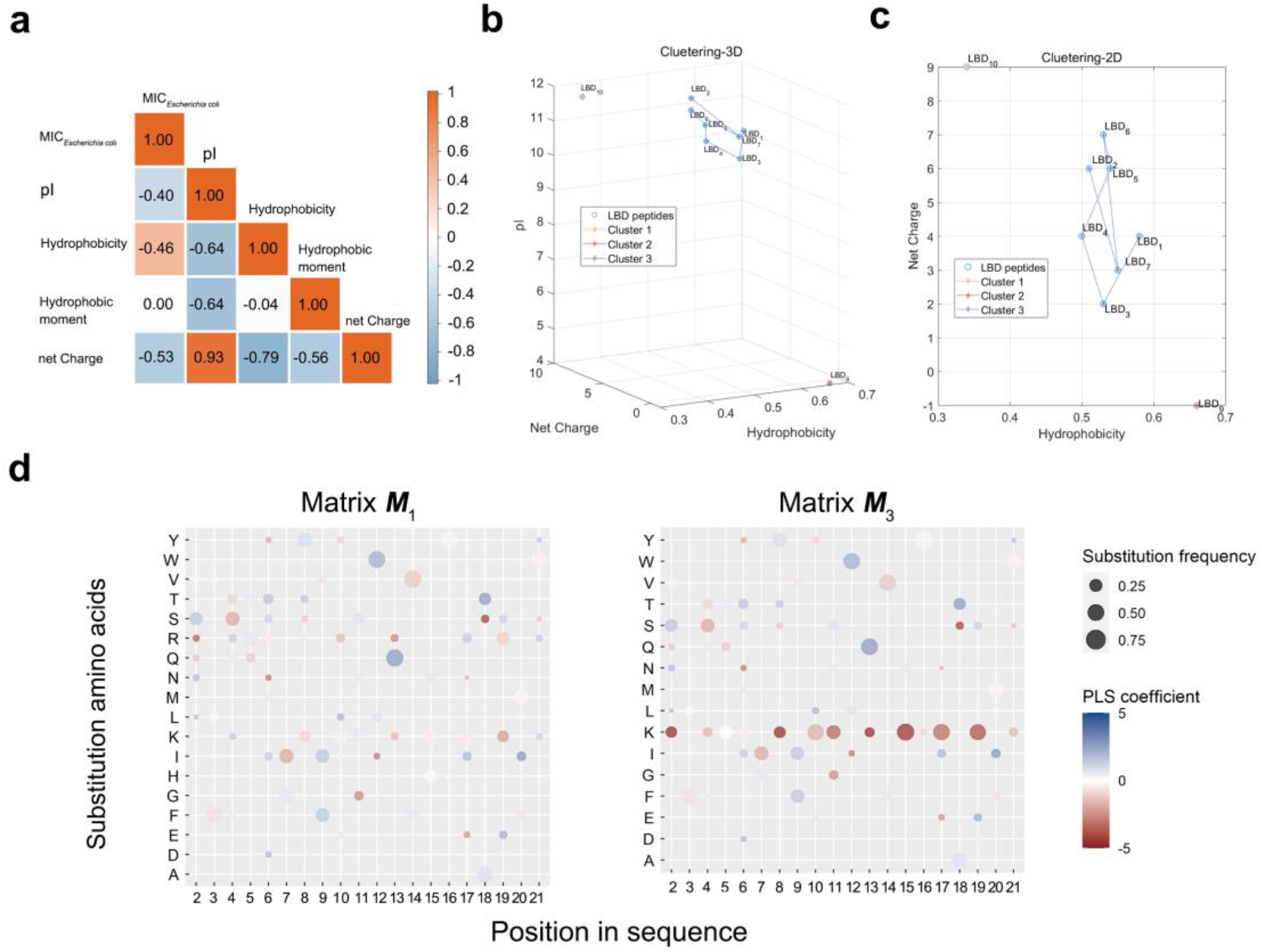
The results from physicochemical property investigation and fitness matrix construction processes. **a** The Pearson correlation coefficient measurement indicates that the MICs and physicochemical properties are not simply in a linearized relationship. **b** and **c** FCM indicate that LBD_1-8_ were the largest cluster created in space of hydrophobicity ranging from 0.5 to 0.7, net charge of 2 to 7, and isoelectric point of 10 to 12. **d** Results of the initial fitness matrix ***M***_1_ generated by PLSR and its derived final fitness matrix ***M***_3_.

In the relative-MIC prediction process, we assigned each optimized LBD sequence in set **K**_2_ with a relative-MIC based on the Euclidean distance of physicochemical properties (net charge and hydrophobicity) between each optimized LBD sequence and LBD_1-8_. This turned out that each optimized LBD sequence in set **K**_2_ was assigned a relative-MIC ranging from 10.0 to 36.7, and approximate 44 optimized LBD sequences have a relative-MIC value ≤ 15.

During iterative exploration, GA was used to improve the population of LBD sequences with relative-MIC ≤ 15, where each iteration was evaluated by the relative-MIC as a fitness function. Among one hundred-round simulation, the average number of iterations was 36, and at this point, the population size of optimized LBD sequences with the relative-MIC ≤ 15 could reach a stationary state (Supplementary Fig. 2). The final 68 optimized LBD sequences were collected from the intersection of each simulation output and performed with the substitution frequency investigation. As shown in Fig. 3a, the substitution of E^13^ and D^8^ with K was encountered most frequently, followed by substitution of N^4^ and Q^10^ with K. Thus, we replaced other residues with K according to the rankings of substitution frequency (Supplementary Table 3); for example, N^4^ should be replaced with K, and then, a new optimized LBD sequence was generated with 3 mutations. To this end, four sequences, LBD_A-D_, were generated and subsequently synthesized for downstream application (Fig. 3b). In addition, two randomly mutated sequences, LBD_X_ and LBD_Y_, were designed as control following the empirical method of simply increasing net charge.

**Fig. 3.**
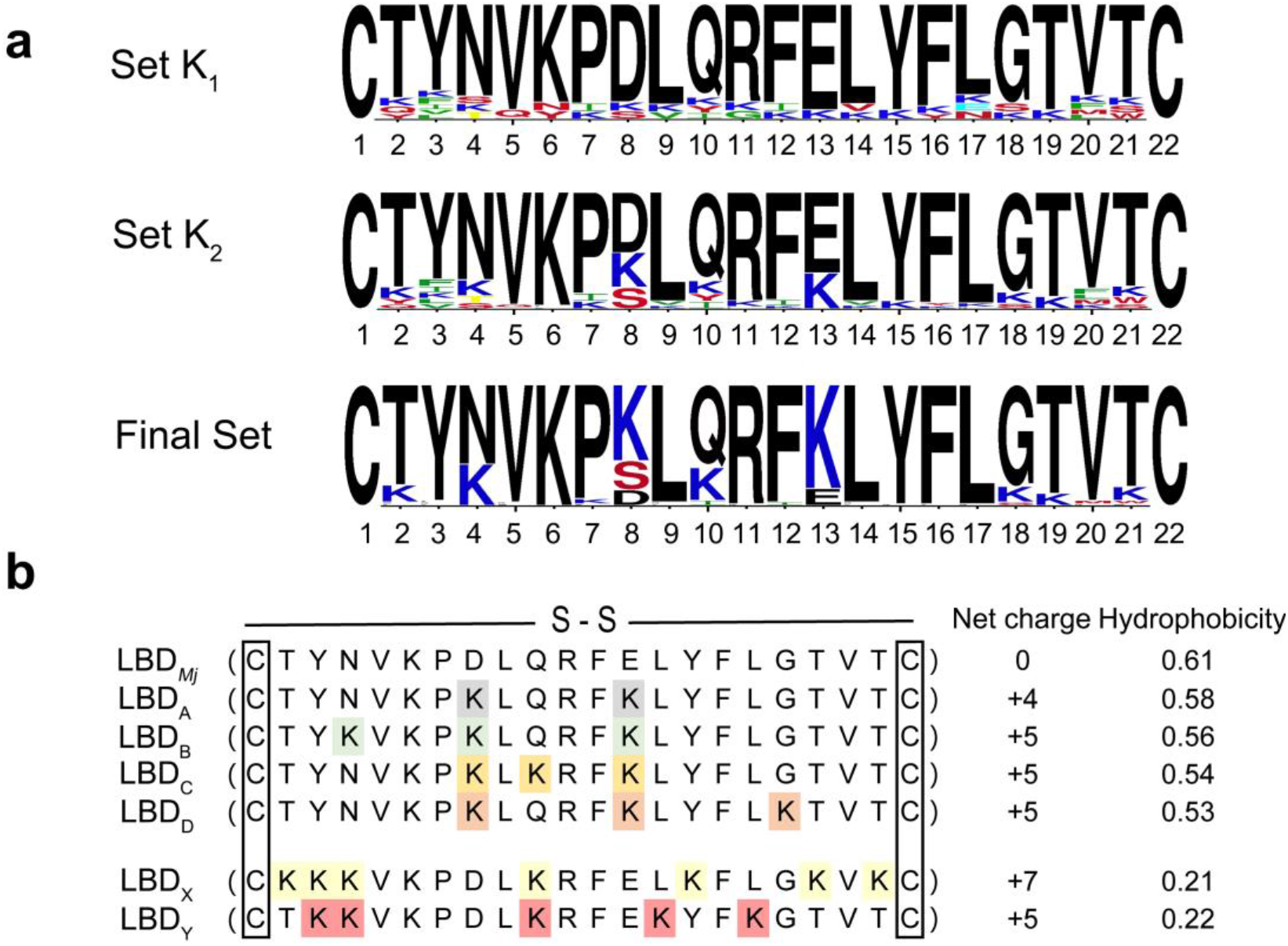
Identification of the substitution sites of amino acids that have strong effect on enhancing anti-*E. coli* ability. **a** Machine learning combining with genetic algorithm initially identifies 9432 newly generated LBD sequences (Set **K**_1_) which was further filtered by retaining only the preset net charge and hydrophobicity, resulting in a new library of optimized LBD sequences (Set **K**_2_) with a reduced number to 1593; **K**_2_ was used for relative-MIC assignment and for the next simulation step, and eventually generating approximate 68 optimized LBD sequences; amino acids from original and modified peptides are indicated by black and color letters, respectively. **b** The sequences of the synthesized peptides, including computing output LBD_A-D_, and two randomly mutated LBD_X_ and LBD_Y_ as controls.

### Antibacterial activity

We evaluated the antibacterial efficacy of the synthetic peptides LBD_A-D_, LBD_X_ and LBD_Y_ along with the original peptide LBD_*Mj*_ by determining their MICs against *E. coli*, plus various gram-negative and gram-positive bacteria (Table 1). The MIC used in this study is defined as the lowest concentration of peptides that inhibits the visible growth of the tested strains within 24 h. Overall, the synthetic peptides LBD_A-D_, LBD_X_ and LBD_Y_ exhibit better antimicrobial activities than does LBD_*Mj*_, as indicated by at least an order of magnitude lower MIC. Also, LBD_A-D_ exhibited much higher antimicrobial (*E. coli)* activity than two randomly synthesized LBD_X-Y_, as indicated by the MIC values of < 2.5 μM and > 40 μM, respectively. Additionally, LBD_X_ and LBD_Y_ showed MICs higher than 40 μM against most of the gram-negative species, even though their net charge value was up to 5-7.

**Table 1.**
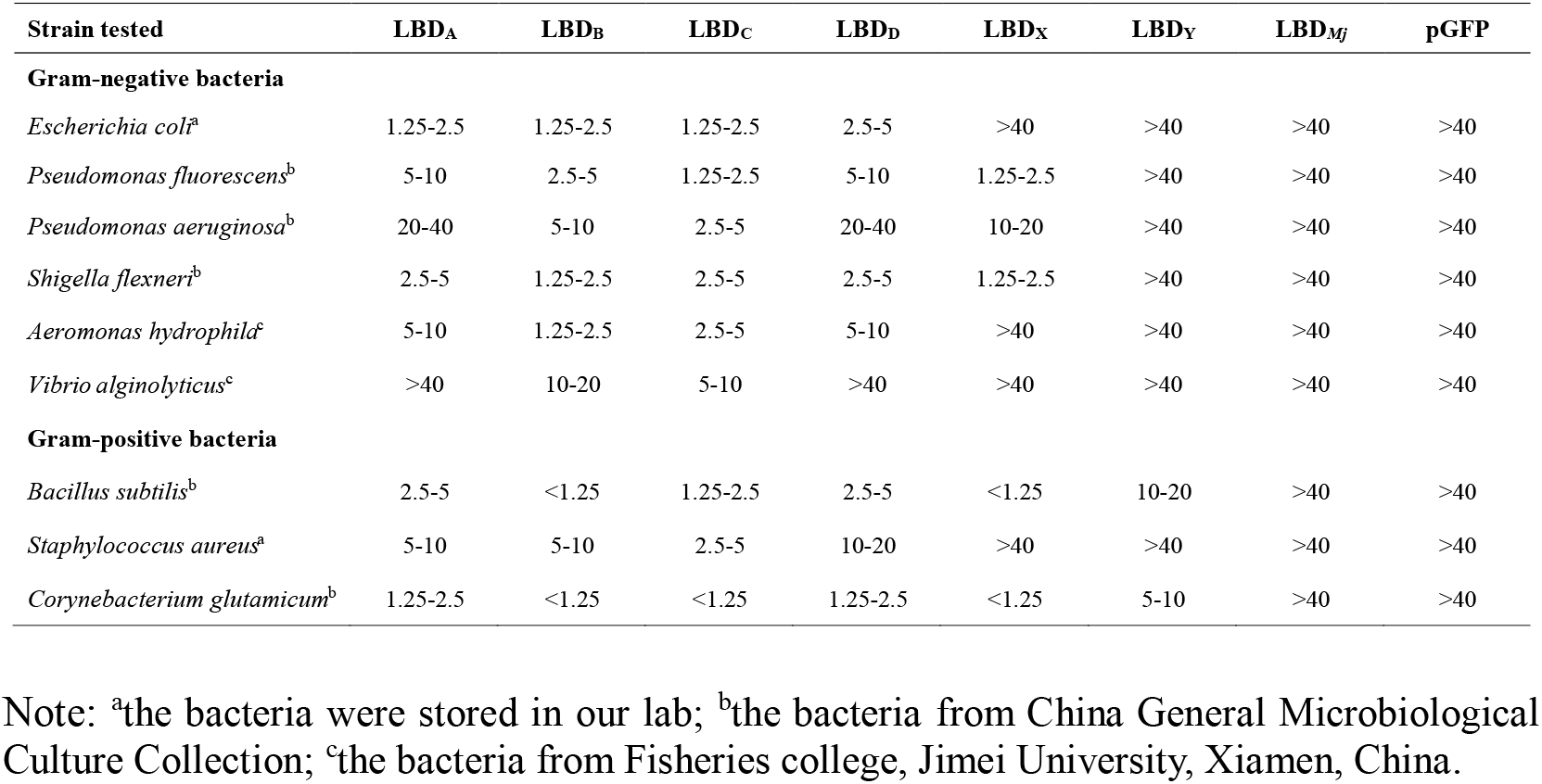
MIC analysis of LBD_A-D_, LBD_X_, LBD_Y_, LBD_*Mj*_, and negative control pGFP peptides (μM).

LBD_B_ showed higher antimicrobial activity than LBD_A, C-D_, LBD_X_ and LBD_Y_, and the highest activity (< 2.5 μM) from LBD_B_ was observed for the gram-negative bacterium *E. coli, Shigella flexnerib, Aeromonas hydrophilac* and the gram-positive bacteria *B. subtilis* and *C. glutamicum.* Again, in our previous study(*36*), 40 μM MIC was chosen as a threshold, and any values higher than that are considered not active; thus, no further experiments regarding bioassays were continued. Overall, the antibacterial spectrum of LBD_A-D_ was broadened compared to that of LBD_*Mj*_, and more interestingly, LBD_A-D_ did not develop any resistance to all strains tested in this study, regardless of whether they were tested at either sub-MIC or sur-MIC levels, at least under our laboratory conditions.

Given that LBD_B_ has the broadest antibacterial spectrum, we then used it for the studies of pharmacodynamics (PD) and pharmacokinetics (PK). Unless otherwise stated, all bioactivity tests were repeated at least three times. Kinetics analyses were performed and shown in Fig. 4a and 4g. Although the killing index of LBD_B_ for *S. aureus* increased over time, it showed much stronger microbicidal efficacy when using 10 or 20 μM for *E. coli*. Most bacteria were killed after 2 h, suggesting time-dependent activity. Moreover, after incubation with gradient LBD_B_ at 37°C for 2 h, the dose-dependent antibacterial activity of LBD_B_ against *E. coli* and *S. aureus* was detected by characterizing the positively correlated mortality against the concentrations (Fig. 4b and 4h). Results showed that LBD_B_ at 1.25 μM killed 73.1% of *E. coli* and 41.1% of *S. aureus* and when the concentration was doubled (2.5 μM), 100% of *E. coli* and 38.7% of *S. aureus* were killed, suggesting that LBD_B_ is particularly effective against gram-negative bacteria.

**Fig. 4.**
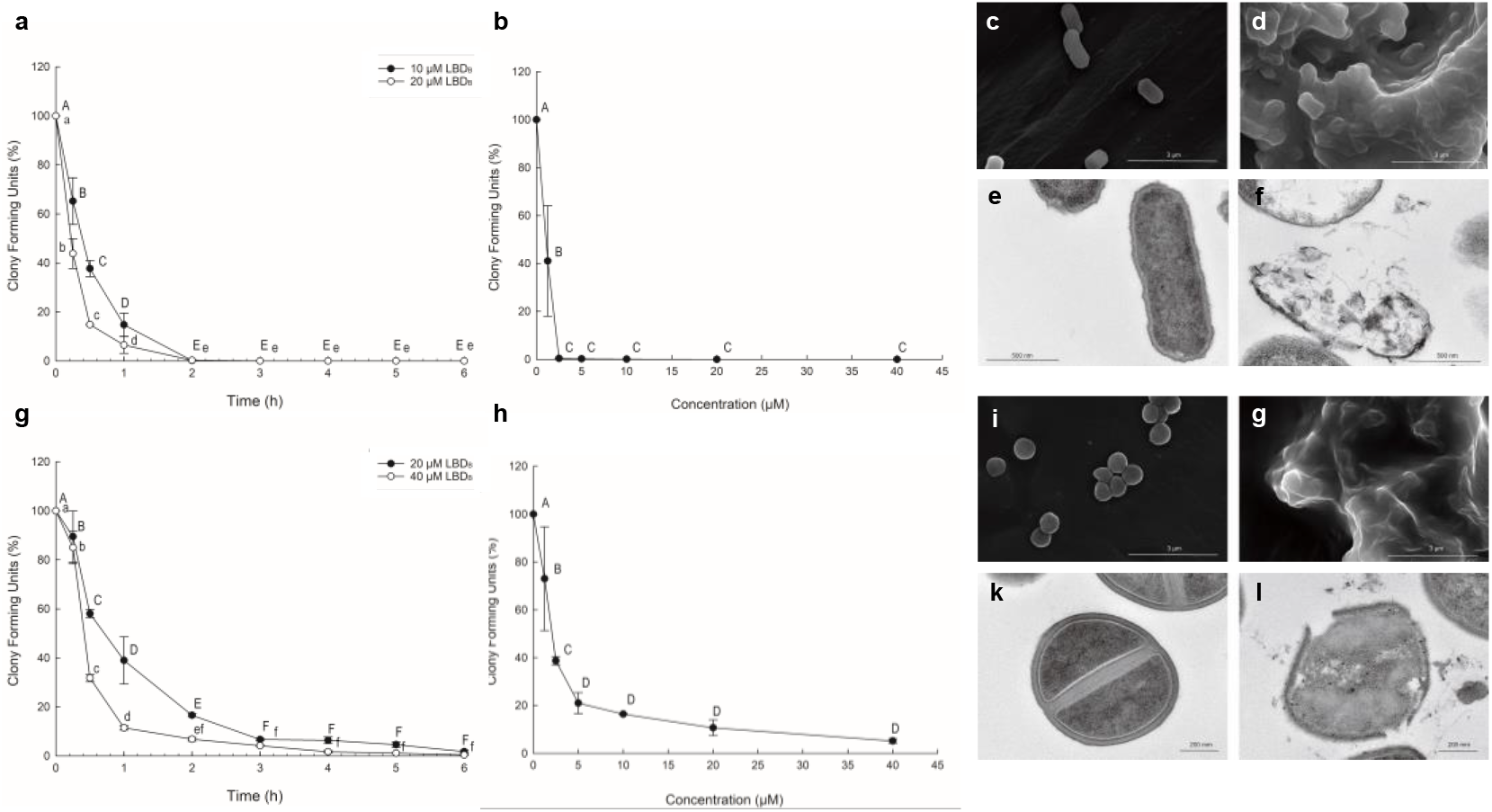
Pharmacokinetics, pharmacodynamics and morphological studies for LBD_B_. PK and PD showing that LBD_B_ exhibiting activities against *E. coli* (**a** and **b**) and *S. aureus* (**g** and **h**) in time dependent (**a** and **g**) and dose dependent (**b** and **h**) manners, respectively; SEM (**c**, **d**, **i** and **g**) showing that the cells of *E. coli* (**d**) and *S. aureus* (**g**) were severely damaged after LBD_B_ treatment due to cytoplasm leakage in comparison with controls (**c** and **i**); TEM showing that a series of pathological changes were observed on the cell surface of *E. coli* (**f**) and *S. aureus* (**l**) with LBD_B_ treatment compared with control groups (**e** and **k**). The same capital letter represents no significant difference between the two groups, whereas same letters in both upper- and lowercase at one data point (time or concentration) represent significant difference between the two groups where uppercase and lowercase letters indicate *p* <0.01 and *p* <0.05, respectively.

### Morphological studies

The antibacterial mechanism was characterized using scanning electron microscopy (SEM) and transmission electron microscopy (TEM) to examine the ultrastructural changes in the tested strains induced by LBD_B_. As shown in Fig. 4, all bacterial cells in the blank (Fig. 4c and 4i) were well-shaped and smooth with a normal form and were not attached. However, the surfaces of *E. coli* and *S. aureus* treated with LBD_B_ were severely damaged due to cytoplasm leakage (Fig. 4d and 4g). Consistent to the above kinetic studies and compared to the control groups, close observation revealed a viscous material around the bacterial cells when the incubation time was prolonged to 2 h after treatment with LBD_B_, and the cell surface became markedly rough and was disrupted with cell debris (Fig. 4d and 4g).

To investigate the detailed microstructure of the cell’s surface, TEM analysis was used to explore the morphological and structural changes of *E. coli* and *S. aureus* cells after incubation with LBD_B_. TEM clearly showed differences in morphology between the untreated and LBDB-treated bacteria. TEM images of bacterial cells from the control groups revealed that cells had a normal cell shape and an inner membrane with an undamaged and intact architecture, and the outer membrane was round and smooth (Fig. 4e and k). In comparison, a series of pathological changes, such as plasmolysis, vacuolation and cytoplasm disruption, occurred in the cells immediately after treatment with LBD_B_ (Fig. 4f and 4l). At 2 h, bacterial cell lysis and content leakage were clearly observed with LBDB treatment (Fig. 4f and 4l). These results indicated that the membrane was permeabilized by LBDB, which caused cell lysis and eventual death.

### NMR solution structure of LBD_B_

To describe the structure and activity relationship, we performed a set of 1D and 2D nuclear magnetic resonance (NMR) experiments for LBD_B_ (Fig. 5). All chemical shifts of protons and carbons from the primary chain and side chains were assigned, except those on the aromatic residues. The refinement statistics for the lowest-energy structure are summarized in Table 2. ^1^H-^1^H NOESY indicated a total number of 291 NMR distances and dihedral restraints with an RMSD of the average pairwise distance of the backbone atoms of 0.74 Å, indicating an average of 12 unambiguous NOEs per residue. The 3D structure of LBD_B_ was calculated using the interproton distance restraints derived from the NOE experiment and the dihedral angle (φ, ψ) restraints based on the chemical shifts. Fig. 5 shows a structural ensemble of 20 representative structures, a representative ribbon structure and an electrostatic potential surface, respectively. The NOE experiment and chemical shift of C_β_ indicated that LBDB possesses an overall extended and rigid structure with a disulfide bond formed between residues C^1^ and C^22^ and two 310-helices (residues P^7^-K^8^-L^9^ and F^12^-K^13^-L^14^, 0.27 Å of backbone RMSD). This conformation is probably caused by the high number of positive charges and high hydrophobicity, and thus consists of no apparent α-helices and β-strands. As shown in Fig. 5, the electrostatic potential map suggests that most of the area of the molecule is highly hydrophobic, as indicated by the hydrophobic N- and C-terminal residues of Y^3^, V^5^, Y^15^, F^16^, L^17^ and V^20^ and the other few residues in the middle sequence, such as L^9^, F^12^, and L^14^. However, there are some polar residues in the middle area that are positively charged (e.g., K^4^, K^6^, K^8^ and R^11^). Therefore, the overall solution structure adopts a partly amphipathic conformation, where the hydrophilic sides were in direct contact with the polar solvent, such as water in this case. In addition, C^1^-T^2^-Y^3^-K^4^-V^5^-K^6^ and F^16^-L^17^-G^18^ are two primary short sequences that fold the entire structure of topological space. Finally, the partial sequence of P^7^-K^8^-L^9^-Q^10^-R^11^-F^12^-K^13^-L^14^-Y^15^ is twisted, while that of T^19^-V^20^-T^21^-C^22^ is markedly kept in a linear conformation.

**Fig. 5.**
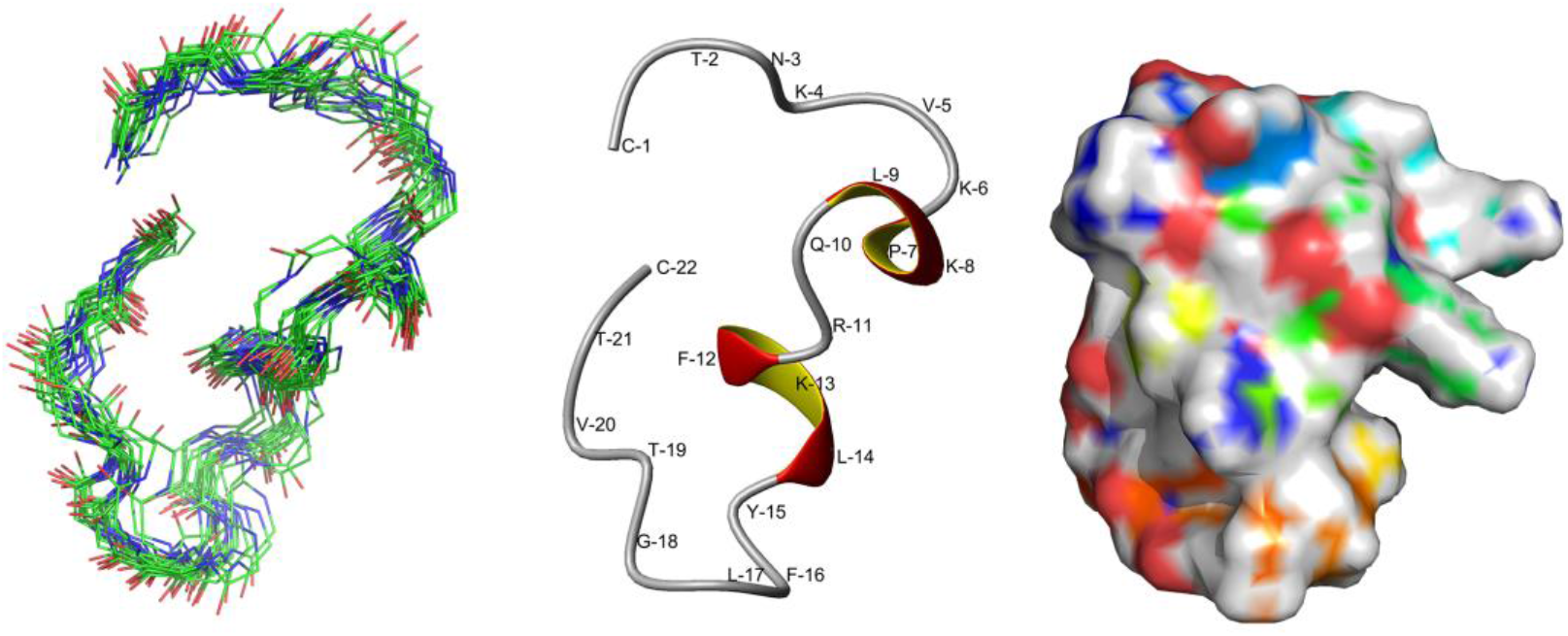
3D solution NMR structure of LBD_B_. Cartoon representation showing: overlaid 20-lowest energy structures (left), a Ribbon extended structure with two 310-helix (middle, disulfide bond not shown), and electrostatic surface representation (right).

**Table 2.** Solution NMR and refinement statistics for LBD_B_ determination.

|  | LBD<sub>B</sub> |
| --- | --- |
| NMR distance and dihedral constraints |  |
| Intra-residue unambiguous NOEs | 141 |
| Sequential unambiguous NOEs | 88 |
| Medium range unambiguous NOEs | 26 |
| Long range unambiguous NOEs | 36 |
| Total unambiguous NOEs | 291 |
| Total ambiguous NOEs | 282 |
| Average pairwise r.m.s. deviation (Å) |  |
| Average backbone atoms | 0.74±0.15 |
| Average heavy atoms | 1.21±0.19 |
| Energy (kcal/mol) |  |
| Mean amber energy | -660.74±3.30 |
| NOE distance restraints violation energy | 3.79±0.66 |
| Torsion angle restraints violation energy | 0.53±0.32 |
| Restraint violations |  |
| Distance (> 0.3 Å) | 0 |
| Dihedral angle (>5°) | 0 |
| Ramachandran statistics (%) |  |
| Residues in most favored regions | 55.8 |
| Residues in additional allowed regions | 36.8 |
| Residues in generously allowed regions | 7.2 |
| Residues in disallowed regions | 0.2 |

Intrigued by the unique structure of LBD_B_ and its promising *in vitro* observations from the morphological studies, we then evaluated the efficacy and safety of LBD_B_ by *in vivo* antibacterial and antiviral experiments.

### *In vivo* efficacy of LBD_B_

The efficacy of LBD_B_ *in vivo* was evaluated by a shrimp lethality assay, and *V. alginolyticus* and *S. aureus* were used as the representatives of gram-negative and gram-positive bacteria, respectively. For the experiments, we co-injected pathogens with LBD_B_, pGFP and sterile saline, along with pure sterile saline as a negative control, into *M. japonicus*. As shown in Fig. 6a and 6b, the cumulative mortality of the experimental group incubated with the pathogen and LBD_B_ was markedly lower than that of the control groups incubated with either saline or pGFP. *V. alginolyticus* had a strong lethal effect on shrimp, as shown in the control groups, but LBD_B_ reduced mortality to approximately 30%, suggesting a good protective role *in vivo* Fig. 6a. The death curve of *S. aureus* shrimp was marginally later than that of the *V. alginolyticus* group. Fig. 6b shows that *S. aureus* also had a strong lethal effect on shrimp but was slightly weaker than *V. alginolyticus*. This observation is also consistent with the proposed conclusion regarding the kinetic analysis in which LBD_B_ shows stronger efficacies against gram-negative pathogens.

**Fig. 6.**
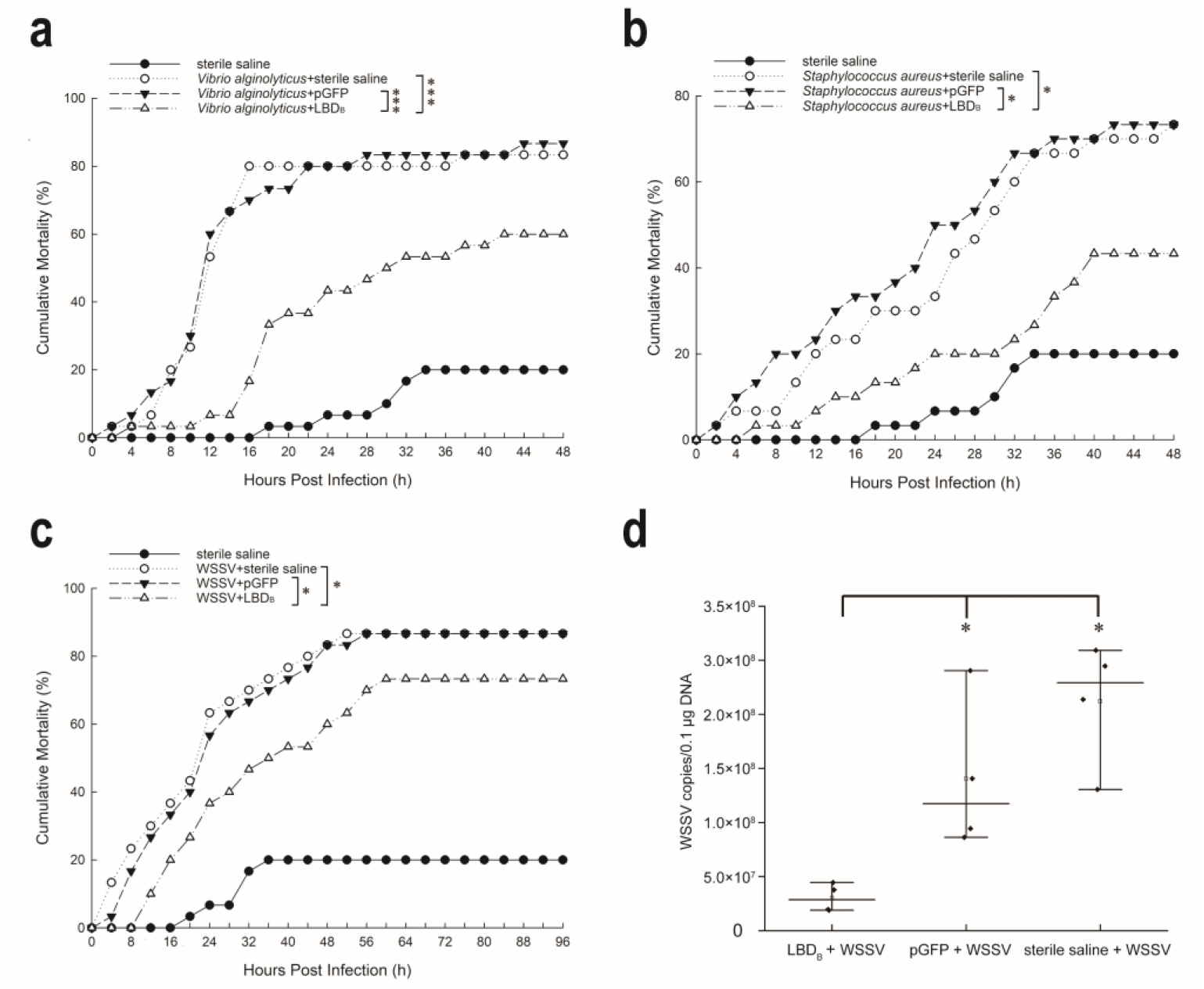
*In vivo* bioactivity test for LBD_B_. Co-injection of LBD_B_ with *V. alginolyticus* (**a**), *S. aureus* (**b**) and WSSV (**c**) in *M. japonicus* showing that LBD_B_ can reduced the mortalities in a reasonable level; the absolute quantitative detection of the virus copy number showing that ‘LBD_B_+WSSV’ group was significantly reduced to 4- and 10-fold in comparison to the ‘pGFP+WSSV’ negative control group and ‘sterile saline-WSSV’ blank control group (*p* < 0.05), respectively (**d**). * indicates *p* < 0.05 and *** indicates *p* < 0.001, respectively.

Similarly, we performed virus infection experiments by co-incubating LBD_B_ with WSSV virus solution in *M. japonicus* at room temperature. Fig. 6c shows individual death in each group at the initial stage after the injection of WSSV, which may be due to stress reactions. Similar to the results of the bacterial clearance experiment, the cumulative mortality of the experimental group incubated with WSSV and LBD_B_ was lower than that of the control groups. Fig. 6d shows the results of absolute quantitative detection in which the virus copy number from ‘LBD_B_+WSSV’ group was significantly reduced to 4- and 10-fold in comparison to the ‘pGFP+WSSV’ negative control group and ‘sterile saline-WSSV’ blank control group (*p* < 0.05), respectively.

## Discussion

The resistance of microorganisms to antibiotics is a severe problem that affects public health and clinical practice. AMPs are emerging alternatives to natural product antibiotics and have been extensively studied due to their vast natural abundance and ability to kill microbes(*43, 44*). Given the antibacterial activity of AMPs is primarily determined by their physicochemical properties but not absolutely depending on their specific amino acid sequences(*45*), repurposing and modifying known natural AMPs may contribute to the successful development of new therapeutic strategies. Well-exemplified studies include the basic amino acid alteration in the LBD sequence of ALF, a major group of crustacean AMPs, which markedly improved antibacterial activity^(*35, 46–48*)^. However, these studies did not show the very clear logics that could indicate which amino acid in the sequence should be altered. As it has been speculated that the appropriate insertion, deletion, substitution, and chemical modification of amino acids could alter the physicochemical properties of natural AMPs, thereby improving antimicrobial activity and reducing mammalian cell cytotoxicity(*42, 47*), the key question then arises as to how to make the decision to precisely alter the peptide sequence to obtain the best activity. In this study, we are guided to use an integrated ensemble machine learning method to evaluate the short LBD sequences generated by directed evolution and build a low-cost but highly reliable transformation algorithm, eventually allowing the minimization of old sequence variation and the maximization of new peptide activity.

The algorithm shown in this work is based on a probabilistic model that performs the directed evolution of LBD_*Mj*_. The key purpose is to find the best candidates of amino acid substitution sites so that the antibacterial activity of the modified peptide with identical length would be enhanced. Because there are only 30 LBD datasets, it is unrealistic to use the currently popular deep learning model such as neural network for data training. In addition, simple alignment of LBD_1-10_ did not show any significant identities or similarities (Supplementary Table 1). However, previous studies revealed the applications of linear model were capable of identifying the advantageous and disadvantageous of the substitution sites of certain peptide sequences(*37, 49*). Therefore, in this work, considering the number of samples was less than that of variables (30 vs 85) when the amino acid replacement matrix (***M***_1_) was constructed, we applied PLSR to linearize the substitution sites of the published peptide sequences and their MICs and at the meantime, used Monte Carlo simulations to enhance the possibility of the substituted positively charged amino acids. This process resulted in a roulette-like fitness matrix in which each coefficient represents the contribution to MIC from the substitution sites where the original peptide was modified (Fig. 2d). Next, we clustered the interesting physicochemical properties, such as the net charge and hydrophobicity, to obtain the chemical space for the substitution sites of protein sequences(*49*), which is consistent with some current studies in which net charge and hydrophobicity were observed to be crucial for the activity of LBD family peptides(*50*).

In this way, the characteristic intervals of the physicochemical properties from LBD_1-8_ sequences were clustered and used as standards for the new peptides generated from the fitness matrix (***M***_3_). Briefly, ***M***_3_ provides the most likely substitution sites of the original peptide, on which the combination of the substitution could generate the optimized peptide sequence that should be within a reasonable interval of physicochemical properties. Although we grouped the MICs of the LBD peptides according to their published results to best avoid the influence of the real value, there may still be errors that occurred in different laboratories. However, based on the proposed model, the MIC value (> or < 30 μM) of the LBD peptide from these two published groups is relatively accurate. The reason that all strong substitution sites are lysine is attributed to the result of merging the distribution coefficients of arginine and histidine into lysine in the dataset. This process is also referred to previous publications in which the parameters were adjusted to best contribute to the antibacterial activity(*42, 44*). It is worth noting that the proposed algorithm accurately substituted two negatively charged amino acids with positively charged amino acids, which increased the net charge without affecting the hydrophobicity, because the hydrophobicity of the input peptide LBD_*Mj*_ itself is within a reasonable range and the only shortcoming of this peptide is the insufficient net charge. We therefore conclude that it is this feature that makes LBD_*Mj*_ difficult to approach *E. coli*, which has a negatively charged phospholipid membrane.

To verify our method, we carried a series of downstream experiments such as total synthesis, structure analysis, *in vitro* and *in vivo* bioassays. Result showed that the antibacterial activity of all synthesized LBD_A-D_ was improved compared to that of LBD_*Mj*_. LBD_A-D_ exhibited stronger *in vitro* antibacterial activity against most strains tested in this study, including *S. aureus* and *V. alginolyticus*. This result is in agreement with that LBD_A-D_ had a more positive net charge and higher hydrophilicity than LBD_*Mj*_. In addition, because amphipathicity with the positively charged polar face and the nonpolar face is required for insertion into the membrane interface as it causes increased permeability and loss of barrier function of target cells(*51*), the rational distribution of hydrophilic and hydrophobic residues in LBD_A-D_ may explain the enhanced antimicrobial activity, as indicated by the NMR solution structure of LBD_B_.

Despite many natural and modified AMPs have demonstrated considerable advantages for antimicrobial applications, some boundaries for drug exploitation exist; and this is mostly because AMPs have low stability *in vivo* due to their complex surrounding environments. However, the engineered LBD_B_ peptide with a short sequence in this study exhibited apparent *in vivo* antibacterial and antiviral activities and thus played a protective role in shrimp. Additionally, a markedly reduced mortality rate was observed in *M. japonicus*, and the copy number of WSSV was significantly lower than that in the control groups. *V. alginolyticus* and WSSV are pathogenic agents that cause massive mortality in the shrimp industry, and the inhibition of pathogenic microbes *in vivo* using the LBD_B_ peptide may provide new strategies for shrimp disease control.

To conclude, we have developed a bespoke algorithm based on feature engineering, machine learning, and genetic algorithms to minimize amino acid replacement of peptide sequences while maximizing antibacterial activity. Importantly, the combination of machine learning and genetic algorithms allows the high accuracy of outputs and reduction of verification cost for the wet experiments. In addition, the results generated by this algorithm could provide valuable guidance for future work such as *in vivo* gene editing so that the modified AMPs will be produced in the host and reduce the possible consequences of the enhancement of haemolytic toxicity. Although some prior knowledge is needed in this algorithm, the concept here is suitable for any AMP modifications and widely applicable. This, fortunately, means the original peptide sequence features mined by the algorithm and the structural features after sequence modification can help to enhance the robustness for directed evolution in the future. In short, our work allows for the rapid design of activity-enhanced peptide sequences with a relatively short chain length and, therefore, opens a new route to generate peptide antibiotics.

## Materials and methods

### LBD sequence acquisition, comparison and physicochemical property calculation

A 22-mer peptide, LBD_*Mj*_ (CTYNVKPDLQRFELYFLGTVTC), was identified from *M. japonicus.* From the known LBD-associated papers, 30 published LBD_1-30_ sequences and their MICs against *E. coli* were collected and are shown in Supplementary Table 1. LBD_1-10_ has a MIC below 30 μM, whereas LBD_11-30_ has a MIC above 30 μM. To normalize the MIC of LBD_1-30_, we defined the mean values of MICs in this manner: the MICs of each published peptide were rearranged in an interval of 10 μM. For example, LBD_1_ has its MIC in between 0 and 10, then, its mean MIC was assigned to 5. The four physicochemical parameters of LBD_1-30_ (i.e., net charge, hydrophobicity, isoelectric point and hydrophobic moment) were calculated using the formula as described in previous work(*52–54*). Multiple sequence alignments were performed using ClustalW(*55*) and are shown using the ESPRIPT 3.0 tool (http://espript.ibcp.fr/).

### Fitness matrix constructions

A substitution matrix (***X***) was generated based on the relative frequencies of amino acids after performing the multiple sequence alignments between LBD_*Mj*_ and LBD_1-30_. The MICs of each corresponding sequence were converted to a matrix (***Y***). The initial fitness matrix (*M*_0_ = (*a_ij_*)_20×20_ was generated from ***X*** and ***Y*** using PLSR, which can forecast a potential increase in the antimicrobial activity of the peptide when an amino acid residue is replaced with another residue. The resulting linear model can be expressed as:

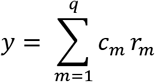

where *c_m_* is the contribution to fit the substitution of each amino acid, and *r_m_* indicates the presence (*r_m_* = 1) or absence (*r_m_*= 0) of the substitution. The obtained PLSR coefficients were used as fitness values in the initial fitness matrix ***M***_0_ and as a probability to introduce substitutions of the amino acids at each round of optimization. Based on the PLSR coefficients, substitutions were categorized as advantageous, detrimental, or innocuous to determine whether they should be kept, eliminated, or retested. The substitution coefficients of K, R and H in each position were merged into one row and all represented by K, and a matrix ***M***_1_ = (*b_ij_*)_18×20_ in this way was degenerately generated.

The positions rather than the number of positively charged residues have been shown to affect biophysical properties, and moreover, the selectivity of K in the peptide more strongly affect antimicrobial activity(*56, 57*). Therefore, the frequency of distribution of positively charged amino acids such as K, R and H in LBD_1-10_ were computed, and a matrix that contains frequency of class intervals of positively charged amino acids ***M***_2_ = (*c*_1*j*_)_1×*t*_ was generated, where *t* is the maximum index interval of positively charged amino acids. For example, when positively charged amino acids appeared in the first and ninth positions, *t* will be denoted to 8. The set of negative values {*b*_18*k*_}, *k* ∈ *K* in the set of substitution coefficients of K was used to reassign the PLSR coefficients in {*b*_18*j*_} according to the ***M***_2_ = (*c*_1*j*_)_1×*t*_. For the *i* position of (*b*_18*j*_), the process of reassignment to this position can be expressed as:

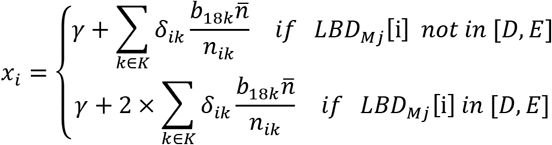

Where 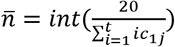 represents the expected number of positions that could be allocated; *n_k_* = *card*(*G_k_*) represents the actual number of positions that could be allocated; *G_k_* is a set of the actual position that could be allocated; *δ_ik_* is a variable indicating 0 or 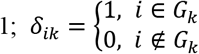; where {*D,E*} is the set of negatively charged amino acids. Details of allocation process is referred to pseudo-code in Supplementary Table 4. Monte Carlo simulation was conducted in which the above allocation process was repeated 3×10^4^ times to derive the probability distributions of the substitution coefficients of positively charged amino acids (Supplementary Fig. 2). The mean values of the coefficient in the range of the highest probability density in each position were reassigned to (*b*_18*j*_). To sum up, ***M***_3_ = (*d_ij_*)_18×20_ were prepared from the matrix ***M***_1_ and ***M***_2_. The initial optimized peptide Set **K**_1_ was generated according to the corresponding fitness values in matrix ***M***_3_ using the model of roulette wheel selection when only negative values in ***M***_3_ were considered.

### Relative-MIC assignment of *in silico* LBD sequence

The Pearson correlation coefficient between MIC values and four physicochemical properties was computed using the corr() in MATLAB R2021b, and the properties with a coefficient with approximate 0.5 were used for subsequent computation. The MIC value, net charge, hydrophobicity and isoelectric point of LBD_1-10_ were clustered using FCM embedded in FCM() function in MATLAB R2021b. The range of net charge and hydrophobicity for the largest cluster was estimated and considered to be a preset range. The secondary optimized sequences in Set **K**_2_ were retained after the filtration from preset range.

The above analysis indicates net charge and hydrophobicity are speculated as two important factors affecting the antimicrobial activity. The similar idea to the K-Nearest Neighbor (KNN) algorithm that we proposed here is to use the distance of net charge and hydrophobicity between the LBD_1-8_ and optimized LBD sequences in set **K**_2_ as a basis to weigh the relative-MIC value for each optimized LBD sequence. The relative-MIC assignment formula is expressed as:

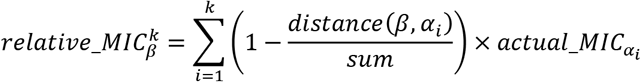

Where *k* is the best value returned by identical assignment process performed on the LBD_1-30_ sequences; 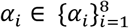 represents any of an arbitrary individual from LBD_1-8_; *distance* between optimized sequences and *α_i_* were calculated by two dimensional features including net charge and hydrophobicity. Similar to the KNN algorithm, choice of an appropriate value for *k* has effect on the reassignment of relative-MICs for the optimized LBD sequences (set **K**_2_) from LBD_1-8_. Thus, the best choice of *k* was evaluated by the minimizing sum of squared difference (SSD) between the competitiveness-index of the relative-MICs and the competitiveness-index of the actual MIC while the assignment process was performed among the LBD_1-30_ sequences. The competitiveness-index in pseudo-code was used to evaluate the ranking of LBD sequences based on the relative-MICs. The details of distance calculation and choice of best *k* value is refereed to pseudo-code in Supplementary Table 5.

### Maximizing population of optimized LBD sequences with a genetic algorithm

The optimized LBD sequences in set **K**_1_ were generated *in silico* by introducing up to three random amino acids in the substitution sites, which were biased by the PLSR coefficient value in matrix ***M***_3_. The smaller the coefficient of amino acid substitution is, the greater the probability of substitution occurring, while the other substitutions with higher coefficients are suppressed. The number of iterations on random substitutions was evaluated until the counts of optimized LBD sequences in set **K**_1_ reached the stationary state. The relative-MIC was assigned to each optimized LBD sequence in set **K**_2_ following the above process. To acquire sufficient optimized LBD sequences with relative-MIC ≤ 15, the genetic algorithm was applied to generate the population L_n_ from the population L_n-1_ (start at set **K**_2_), and each round of population was evaluated according to the relative-MICs. For each iteration, those sequence pairs from population L_n_ that have relative-MICs ≤ 15 were chosen to cross with those pairs with relative-MICs ≥ 15. The crossover rate was set to 30% and the generated offspring for population L_n+1_ in which one residue in each offspring was randomly selected for mutation with a chance of 5%. The candidate replacement residues were supplied by a list which included 18 common amino acids (aspartic acid and glutamic acid were excluded due to their negative charge). The iterations were processed until the counts of optimized LBD sequences with a relative-MIC ≤ 15 had been maximized (Supplementary Fig. 3). One hundred independent simulations in this algorithm were performed, and the intersection of optimized LBD sequences of each final output was collected. The advantageous substitution rate was determined by the frequency with which a particular substitution appears in the final set of the optimized LBD sequences with a relative-MIC ? 15. The optimized sequences that had the two highest frequencies of K substitution were identified and further mutated with one residue according to the rank of frequency to generate a new set of optimized sequences. Thus, the sequences generated from the last generalization process were selected for synthesis and *in vitro/vivo* activity tests.

### Synthesis of LBD sequences

To enhance the activity and explore the structure-activity relationships of the LBD_*Mj*_ peptide, four modified peptides, LBD_A-D_, along with LBD_X_ and LBD_Y_, were designed and synthesized based on the analysis of the physicochemical properties and structural information of the LBD of ALF_*Mj*_ (GenBank accession No. MN416688). All synthesized LBDs were protected by leucine at the N-terminus and proline at the C-terminus, which were further acetylated and amidated at each terminus, respectively. Additionally, a disulphide bond was formed between these two cysteine residues (Fig. 3b). The negative control was a partial peptide from green fluorescent protein (GFP) (accession number: AAN41637). All synthetic peptides were commercially synthesized by a biotechnology company (Shanghai Ziyu Biotech Co., Ltd.) and confirmed using mass spectrometry. The purity was tested using high-performance liquid chromatography to ensure a > 95% purity grade. All of the synthetic peptides were dissolved in aseptic deionized water and directly used in antibacterial experiments. The entire process was performed on ice.

### NMR experiment

1D and 2D NMR experiments were performed using a Bruker AVANCE III-700 NMR spectrometer equipped with a cryo-probehead. The LBD_B_ sample was dissolved in 500 μL of H2O with 10% D2O added, resulting in a final concentration of approximately 3.5 mM. The ^1^H-^1^H nuclear Overhauser effect spectroscopy (NOESY) spectra were acquired using a mixing time of 350 ms. All NMR spectra were processed using NMRPipe(*58*) and analyzed by NMRView(*59*). The parameters for LBD_B_ structure calculation include interproton distance restraints derived from nuclear NOE and dihedral angles (φ, ψ) from chemical shifts calculated by TALOS(*60*). The LBD_B_ structure was calculated by CYANA(*67*) and further refined using AMBER(*62*). The CANDID(*63*) function of the CYANA program was used to generate the initial structures, from which 20 lowest energy structures were selected to extend the NOE assignments by SANE(*64*). Then, 100 structures with the lowest energy from a total number of 200 structures calculated by CYANA were selected as the initial structures for refinements using AMBER. Finally, the 20 lowest energy conformers were selected to represent the LBDB structure, which was analyzed using MOLMOL(*65*) and PROCHECK_ NMR(*66*). The NMR data are available in Supplementary Fig. 4-9.

### *Marsupenaeus japonicus* sampling

All healthy prawns (6.75 ± 0.42 g) were collected from Dongshang, Fujian, China and were acclimated in flat-bottomed rectangular tanks with aerated seawater at approximately 23°C and 28% salinity. The water was changed daily for at least one week prior to the infection experiment, and the shrimp were fed twice daily with commercial feed (Fuxing (Xiamen, China) organism feed co. LTD, at least 45% crude protein content).

### Antibacterial test

The antibacterial spectrum detection assay was performed as described previously(*36, 67, 68*). We selected 6 Gram-negative bacteria (*E. coli, Pseudomonas fluorescens, P. aeruginosa, S. flexneri, A. hydrophila* and *V. alginolyticus)* and 3 Gram-positive bacteria (*B. subtilis, S. aureus, C. glutamicum)* for the liquid growth inhibition assay. The assay was performed in 96-well flat-bottom cell culture plates to identify the MICs. All frozen glycerol bacterial stocks were activated via inoculation on lysogeny broth (LB) culture. A monoclonal bacterium of each tested bacterium was chosen for single inoculation on Mueller-Hinton agar (MHA) for culture at 28°C or 37°C. Overnight cultured microbes were diluted to a final OD600 = 0.003 and a final OD600 = 0.006 (marine bacteria). Then, a 2-fold serial dilution of LBDs with final concentrations ranging from 1.25 to 40 μM was incubated with the microbes at 28°C for 24 h.

### Dose-dependent antibacterial activity and time-killing curves

Kinetic studies were performed to further determine the bactericidal rate of LBDB according to previous procedures(*36, 69, 70*), and we chose *E. coli* and *S. aureus* as representative gram-negative and gram-positive bacteria, respectively. Logarithmic phase cultures of *E. coli* and *S. aureus* were incubated with LBDB as described above, and LBDB was replaced with sterile Milli-Q water as a control. For dose-dependent antibacterial activity detection, the bacteria and gradient LBD_B_ were adjusted and cultured for 2 h at 37°C after mixing with 50 μL, and 0.5 μL of the mixture was placed onto MHA and cultured at 37°C for 12-18 h. For time-killing curve detection, 0.5 μL of the mixture was transferred into 50μL of NaPB buffer at various time points (0–360 min). Colonies were counted after plating on MHA plates and incubation at 37°C overnight. The rate of colony forming units (CFUs) was characterized relative to the CFUs obtained within the control (100% CFU at 0 min). The information at each time point is described as the means ± standard deviations (SD) of 3 replicates. One-way examination of change (ANOVA) was used to determine the statistical significance between each group by the SPSS 17.0 program.

### Scanning electron microscopy (SEM) analysis

SEM was used to observe changes in morphology and analyse the antibacterial effects of LBD_B_ according to previously described procedures(*26*). Cultures of *E. coli* and *S. aureus* were incubated with LBD_B_ (final concentration of 40 μM) at 37°C for 15 min, 30 min, 1 h and 2 h. The samples were immobilized with 2.5% glutaraldehyde at 4°C overnight and washed with 1× PBS three times. The samples were dehydrated in a graded series of ethanol solutions for critical point drying, covered with gold and observed and photographed using a FEI Quanta 650FEG scanning electron microscope. Sterile Milli-Q water and pGFP in place of LBD_B_ were used as blank and negative controls, respectively.

### Transmission electron microscopy (TEM) analysis

According to the procedures mentioned above, the collected bacterial cells were treated with 40 μM LBD_B_ for 1 h and 2 h. The effects of LBD_B_ treatment on *E. coli* and *S. aureus* were observed using TEM. Sterile Milli-Q water and pGFP were used as the blank and negative controls, respectively. The bacterial pellets were fixed overnight at 4°C and washed three times with 1×PBS. The bacterial pellets were then transferred to agar embedding blocks and fixed with 2.5% glutaraldehyde in 1×PBS overnight at 4°C. The fixation solution was removed, and the bacteria were soaked in 1× PBS for 15 min at 4°C, 3 times in total. The samples were examined using a Hitachi HT-7800.

### Identification of anti-pathogen activity *in vivo*

To investigate the pathogen resistance of *M. japonicus* and the anti-pathogen activity of LBD_B_ peptide *in vivo*, healthy prawns (n = 40 in each group) were used. The prawns were divided into experimental groups (LBD_B_+*V. alginolyticus*, LBD_B_+*S. aureus* and LBD_B_+WSSV), a blank control group (sterile saline), negative control groups (pGFP+*V. alginolyticus*, pGFP+*S. aureus* and pGFP+WSSV), and positive control groups (sterile saline+*V. alginolyticus* sterile saline+*S. aureus* and sterile saline+WSSV). The synthetic LBD_B_ peptide, with a final concentration of 40 μM, was mixed by shaking and incubated with *V. alginolyticus*, *S. aureus* and WSSV at room temperature for 30 min. 50 μL of the mixture containing 1.25 × 10^6^ CFU bacteria or 1.25×10^6^ copies of WSSV was immediately injected into shrimp in the LBD_B_+*V. alginolyticus*, LBD_B_+*S. aureus* or LBD_B_+WSSV groups. The synthetic pGFP peptide or sterile saline was mixed with bacteria or WSSV and injected into shrimp in the control groups instead of the LBD_B_ peptide. Cumulative mortality was recorded every 4 h. Differences in mortalities between groups were tested for statistical significance using the Kaplan-Meier plot (log-rank X^2^ test) in GraphPad Prism 8 software.

For the WSSV challenge experiments, muscle virus titres sampled from a parallel challenge experiment were detected according to Li *et al.*(*71*). Briefly, muscle tissues from surviving prawns were sampled 96 h post-injection with 4 prawns in each sample, and muscle DNA was extracted using a Marine Animals DNA Kit (TIANGEN, China) according to the user’s manual. The quantities of WSSV genome copies were measured using absolute real-time quantitative PCR with the primers WSSV32678-F/WSSV32753-R and a TaqMan probe (shown in Supplementary Table 6) as described previously(*72*). The WSSV genome copy numbers in 1 μg of shrimp muscle DNA were calculated. The results of synthetic TaqManprobe-WSSV32706 mass spectrometry are shown in the supplemental figure.

## Data availability

The authors declare that the data and materials supporting the findings reported in this study are available upon reasonable request.

## Authors contributions

Z.Q. conceived the original idea. H.Z. and Z.Q. designed the research. H.Z. and Y.Z. performed the biological experiment. J.H., Y.W., P.Y., Y.L. and K.J. performed the bioinformatic and computing analysis. Z.Q. H.Z. and Y.G. analysed the NMR data. H.Z. performed the TEM and SEM experiments. Z.Q., J.H. and H.Z. wrote the paper and all authors reviewed and commented the manuscript.

## Acknowledgments

This work was supported by the National Natural Science Foundation of China (32170079 to Z.Q., 32200035 to H.Z., 12205012 to Y.L. and 12101054 to K.J.), the Natural Science Foundation of Guangdong (2021A1515012026 to Z.Q.), Beijing Normal University via the Youth Talent Strategic Program Project (310432104 to Z.Q) and Startup Project Funding (111032103 to H.Z. and 110321001 to J.H.), the College Youth Innovative Talent Program of Department of Education of Guangdong Province (2022KQNCX155 to H.Z.). Z.Q. thanks Professor Jun Zou from Shanghai Ocean University for his kind comments. H.Z. is grateful to Professor Yong Mao from Xiamen University and Professor Xiangyong Yu from South China Agricultural University for their excellent technical supports, and to Dongshan Swire Marine Station (Xiamen University) for providing the laboratory space. We also thank Dr. Hongwei Li from the NMR facility of National Center for Protein Sciences at Peking University for assistance with peptide structure elucidation.

