## Supplementary Information for "Machine learning-guided directed evolution for the development of small-molecule antibiotics originating from antimicrobial peptides"

**Supplementary Table 1.** General information of $LBD_{1-30}$ originated from ALF, physicochemical properties and MIC-value against *E. coli*.

| **Name** | **Number** | **Sequence** | **Genebank Number** | **Net Charge** | **Hydro- -phobicity** | **pI** | **Hydrophobic moment** | **Measured** $MIC_{E.coli}$ | **Defines Mean** $MIC_{E.coli}$ | **Ref** |
|---|---|---|---|---|---|---|---|---|---|---|
| MnALF4 | $LBD_1$ | CSFQVKPKLRKFQLYFEGKFWC | KR013244 | +4 | 0.58 | 10.91 | 0.10 | 8.5 | 5 | (*1*) |
| SpALF5 | $LBD_2$ | CVLSRYIKLRGLKLKYRATLTC | HF952161 | +6 | 0.51 | 11.74 | 0.04 | 1.69-3.37 | 5 | (*2*) |
| FcALF-LBD1 | $LBD_3$ | CTYSVTPTVKSFELYFKGRMSC | JX853774 | +2 | 0.53 | 10.42 | 0.24 | 16-32 | 25 | (*3*) |
| MrALF | $LBD_4$ | CQYSVNPKIKRFELYFKGRMWC | HQ231235 | +4 | 0.50 | 10.75 | 0.13 | 1.6 | 5 | (*4*) |
| Penmon ALF-B1 | $LBD_5$ | CKFTVKPYLKRFQVYYKGRMWC | ABP73289 | +6 | 0.54 | 10.91 | 0.19 | 5 | 5 | (*5*) |
| Litsty ALF-B1 | $LBD_6$ | CRFTVKPYIKRIQLHYKGKMWC | AGH32549 | +7 | 0.53 | 11.24 | 0.14 | 2.5 | 5 | (*5*) |
| FcALF-LBD4 | $LBD_7$ | CRYSQRPSFYRWELYFNGRMWC | JX853777 | +3 | 0.55 | 10.91 | 0.06 | 8-16 | 15 | (*3*) |
| LvALF8-LBD | $LBD_8$ | CSYSTRPYFLRWRLKFKSKVWC | AVP74305 | +6 | 0.57 | 11.74 | 0.04 | 6 | 5 | (*6*) |
| FcALF-LBD6 | $LBD_9$ | CSYSVKPDIQGFELYFIGSVTC | JX853779 | -1 | 0.66 | 4.05 | 0.14 | 16-32 | 25 | (*3*) |
| SALF24 | $LBD_{10}$ | CHIRRKPKFRKFKLYHEGKFWC | B5TTX7 | +9 | 0.34 | 11.74 | 0.07 | 16-32 | 25 | (*7*) |
| ALFPm11-LBD | $LBD_{11}$ | CSYSTRPYFLRWQLKFKTKIWC | JR205808 | +5 | 0.65 | 11.24 | 0.08 | 128 | 128 | (*8*) |
| FcLBD8 | $LBD_{12}$ | CSYSTRPYFIRWQLKFKTKIWC | MH998632 | +5 | 0.65 | 11.24 | 0.08 | 32-64 | 45 | (*9*) |
| FcLBD8Q | $LBD_{13}$ | CSYSTRPYFQRWQLKFKTKIWC | MH998632 | +5 | 0.56 | 11.24 | 0.11 | 32-64 | 45 | (*9*) |
| FcLBD8G | $LBD_{14}$ | CSYSTRPYFIRWQLKFKGKIWC | MH998632 | +5 | 0.64 | 11.24 | 0.09 | > 64 | 64 | (*9*) |
| Mm-ALF | $LBD_{15}$ | CKFTVKPYIKRFQLNYKGRMWC | MN603756 | +6 | 0.50 | 11.24 | 0.11 | > 40 | 40 | (*10*) |
| FcALF-LBD7 | $LBD_{16}$ | CKFTVKPYIKRFQLYYKGRMWC | AY859500 | +6 | 0.57 | 10.91 | 0.18 | 32-64 | 45 | (*3*) |
| FcALF-LBDr | $LBD_{17}$ | CKFTVKPYIKSFQLYYKGSMWC | AY859500 | +4 | 0.66 | 10.49 | 0.24 | > 64 | 64 | (*11*) |
| FcALF-LBDk | $LBD_{18}$ | CSFTVTPYIQRFQLYYIGRMWC | AY859500 | +2 | 0.83 | 10.58 | 0.07 | > 64 | 64 | (*11*) |
| FcALF-LBD2 | $LBD_{19}$ | CSFNVTPKFKRWQLYFRGRMWC | JX853775 | +5 | 0.60 | 12.23 | 0.12 | > 64 | 64 | (*3*) |
| EcLBD1 | $LBD_{20}$ | CSFQVKPKIKRWQLYFIGTMYC | KY305181 | +4 | 0.69 | 10.91 | 0.16 | > 64 | 64 | (*12*) |
| EcLBD3 | $LBD_{21}$ | CNYRVDPKIKRFQLYFKGRMWC | No mention | +5 | 0.45 | 11.24 | 0.12 | > 64 | 64 | (*13*) |
| EcLBD2 | $LBD_{22}$ | CSYQVKPTIRKFELYFKGTFWC | No mention | +3 | 0.62 | 10.58 | 0.24 | > 64 | 64 | (*13*) |
| LvALF-F | $LBD_{23}$ | CTYFVTPKVKSFELYFKGRMTC | KJ000049 | +3 | 0.57 | 10.58 | 0.21 | > 40 | 40 | (*14*) |
| FcALF-LBD3 | $LBD_{24}$ | CVYSVKPTFQRWQLYFIGSMWC | JX853776 | +2 | 0.85 | 10.58 | 0.16 | > 64 | 64 | (*3*) |
| FcALF-LBDn | $LBD_{25}$ | CKYSQKPSFKRWQLYFKGRMWC | AY859500 | +6 | 0.47 | 11.24 | 0.14 | 32-64 | 45 | (*11*) |
| PtALF8 | $LBD_{26}$ | CNYRVMPRFENWRFYFKGDVWC | KJ081863 | +2 | 0.57 | 10.58 | 0.11 | > 64 | 64 | (*15*) |
| LvALF-E | $LBD_{27}$ | CYVNRSPYLKKFEVHYRADVKC | FE069658 | +4 | 0.33 | 10.42 | 0.20 | > 40 | 40 | (*14*) |

| | | | | | | | | | | |
|---|---|---|---|---|---|---|---|---|---|---|
| FcALF-LBD5 | $LBD_{28}$ | CLLSRSPYLKKLEVHYRAELKC | JX853778 | +4 | 0.43 | 10.58 | 0.20 | > 64 | 64 | (*3*) |
| EcLBD5 | $LBD_{29}$ | CQYKRIPYIKRLELHYRAEVRC | No mention | +5 | 0.36 | 10.75 | 0.15 | > 64 | 64 | (*13*) |
| EcLBD4 | $LBD_{30}$ | CVYKRIGYFYKWELNYKAEVRC | No mention | +3 | 0.47 | 10.25 | 0.21 | > 64 | 64 | (*13*) |

**Supplementary Table 2.** The counts of optimized sequences in each set generated from fitness matrix after 10 times simulation.

| **Output** | **1** | **2** | **3** | **4** | **5** | **6** | **7** | **8** | **9** | **10** |
|---|---|---|---|---|---|---|---|---|---|---|
| **Set K1** | 9467 | 9352 | 9339 | 9501 | 9440 | 9363 | 9435 | 9487 | 9498 | 9445 |
| **Set K2** | 1599 | 1589 | 1582 | 1605 | 1587 | 1585 | 1589 | 1580 | 1604 | 1610 |
| **Target set (relative-MIC, < 15)** | 44 | 44 | 44 | 44 | 44 | 45 | 44 | 44 | 44 | 44 |

**Supplementary table 3.** The frequency (%) of amino acids in the optimized sequences in the final set $\mathbf{K}_3$.

| | **Pos 1** | **Pos 2** | **Pos 3** | **Pos 4** | **Pos 5** | **Pos 6** | **Pos 7** | **Pos 8** | **Pos 9** | **Pos 10** | **Pos 11** | **Pos 12** | **Pos 13** | **Pos 14** | **Pos 15** | **Pos 16** | **Pos 17** | **Pos 18** | **Pos 19** | **Pos 20** |
|---|---|---|---|---|---|---|---|---|---|---|---|---|---|---|---|---|---|---|---|---|
| **G** | 0.0 | 0.0 | 0.0 | 0.0 | 0.0 | 0.0 | 0.0 | 0.0 | 0.0 | 0.0 | 0.0 | 0.0 | 0.0 | 0.0 | 0.0 | 0.0 | 83.8 | 0.0 | 0.0 | 0.0 |
| **A** | 0.0 | 0.0 | 0.0 | 0.0 | 0.0 | 0.0 | 0.0 | 0.0 | 0.0 | 0.0 | 0.0 | 0.0 | 0.0 | 0.0 | 0.0 | 0.0 | 0.0 | 0.0 | 0.0 | 0.0 |
| **V** | 0.0 | 1.5 | 0.0 | 98.5 | 0.0 | 0.0 | 0.0 | 2.9 | 0.0 | 0.0 | 0.0 | 0.0 | 1.5 | 0.0 | 0.0 | 0.0 | 0.0 | 0.0 | 98.5 | 0.0 |
| **L** | 0.0 | 0.0 | 0.0 | 0.0 | 0.0 | 0.0 | 0.0 | 95.6 | 0.0 | 0.0 | 0.0 | 0.0 | 0.0 | 98.5 | 0.0 | 98.5 | 0.0 | 0.0 | 0.0 | 0.0 |
| **I** | 0.0 | 0.0 | 0.0 | 0.0 | 0.0 | 1.5 | 0.0 | 0.0 | 2.9 | 0.0 | 1.5 | 0.0 | 0.0 | 0.0 | 0.0 | 0.0 | 0.0 | 0.0 | 0.0 | 0.0 |
| **P** | 0.0 | 0.0 | 0.0 | 0.0 | 0.0 | 94.1 | 0.0 | 0.0 | 0.0 | 0.0 | 0.0 | 0.0 | 0.0 | 0.0 | 0.0 | 0.0 | 0.0 | 0.0 | 0.0 | 0.0 |
| **F** | 0.0 | 1.5 | 0.0 | 0.0 | 0.0 | 0.0 | 0.0 | 0.0 | 0.0 | 0.0 | 98.5 | 14.7 | 0.0 | 0.0 | 98.5 | 0.0 | 0.0 | 0.0 | 0.0 | 0.0 |
| **Y** | 0.0 | 95.6 | 0.0 | 0.0 | 0.0 | 0.0 | 0.0 | 0.0 | 1.5 | 0.0 | 0.0 | 0.0 | 0.0 | 0.0 | 1.5 | 0.0 | 0.0 | 0.0 | 0.0 | 0.0 |
| **W** | 0.0 | 0.0 | 0.0 | 0.0 | 0.0 | 0.0 | 0.0 | 0.0 | 0.0 | 0.0 | 0.0 | 0.0 | 0.0 | 0.0 | 0.0 | 0.0 | 0.0 | 0.0 | 0.0 | 0.0 |
| **S** | 0.0 | 0.0 | 0.0 | 0.0 | 0.0 | 0.0 | 26.5 | 0.0 | 0.0 | 0.0 | 0.0 | 0.0 | 0.0 | 0.0 | 0.0 | 0.0 | 1.5 | 0.0 | 0.0 | 2.9 |
| **T** | 83.8 | 0.0 | 1.5 | 0.0 | 0.0 | 0.0 | 0.0 | 0.0 | 0.0 | 0.0 | 0.0 | 0.0 | 0.0 | 0.0 | 0.0 | 0.0 | 0.0 | 85.3 | 0.0 | 80.9 |
| **C** | 0.0 | 0.0 | 0.0 | 0.0 | 0.0 | 0.0 | 0.0 | 0.0 | 0.0 | 0.0 | 0.0 | 0.0 | 0.0 | 0.0 | 0.0 | 0.0 | 0.0 | 0.0 | 0.0 | 0.0 |
| **M** | 0.0 | 0.0 | 0.0 | 0.0 | 0.0 | 0.0 | 0.0 | 0.0 | 0.0 | 0.0 | 0.0 | 0.0 | 0.0 | 0.0 | 0.0 | 0.0 | 0.0 | 0.0 | 0.0 | 0.0 |
| **N** | 0.0 | 0.0 | 60.3 | 0.0 | 0.0 | 0.0 | 0.0 | 0.0 | 0.0 | 0.0 | 0.0 | 0.0 | 0.0 | 0.0 | 0.0 | 0.0 | 0.0 | 0.0 | 0.0 | 0.0 |
| **Q** | 2.9 | 0.0 | 0.0 | 1.5 | 0.0 | 0.0 | 0.0 | 0.0 | 69.1 | 0.0 | 0.0 | 0.0 | 0.0 | 0.0 | 0.0 | 0.0 | 0.0 | 0.0 | 0.0 | 0.0 |
| **D** | 0.0 | 0.0 | 0.0 | 0.0 | 0.0 | 0.0 | 13.2 | 0.0 | 0.0 | 0.0 | 0.0 | 0.0 | 0.0 | 0.0 | 0.0 | 0.0 | 0.0 | 0.0 | 0.0 | 0.0 |
| **E** | 0.0 | 0.0 | 0.0 | 0.0 | 0.0 | 0.0 | 0.0 | 0.0 | 0.0 | 0.0 | 0.0 | 0.0 | 97.1 | 0.0 | 0.0 | 0.0 | 0.0 | 0.0 | 0.0 | 0.0 |
| **K** | 13.2 | 1.5 | 38.2 | 0.0 | 100.0 | 4.4 | 60.3 | 1.5 | 26.5 | 0.0 | 0.0 | 85.3 | 1.5 | 1.5 | 0.0 | 1.5 | 14.7 | 14.7 | 1.5 | 16.2 |
| **R** | 0.0 | 0.0 | 0.0 | 0.0 | 0.0 | 0.0 | 0.0 | 0.0 | 0.0 | 100.0 | 0.0 | 0.0 | 0.0 | 0.0 | 0.0 | 0.0 | 0.0 | 0.0 | 0.0 | 0.0 |

**Supplementary Table 4.** Pseudocode of Algorithm 1.

**Algorithm 1** Secondary fitness matrix constructions

*# Acquiring expected number of positions that could be allocated*

**Def** get_n_hat ($M_2 = (c_{1j})_{1\times\theta}$):

$\bar{n}$ = 0

index_interval = 0

**for all** $j = [1, \cdots, t]$ **do**

  index_interval = index_interval+j× $c_{1j}$ *# Acquiring expectation of index intervals*

**end for**

$\bar{n}$ = int(20÷index_interval)

**Return** $\bar{n}$

*# Acquiring expected number of positions that could be allocated*

**Def** positive_charge_index_interval ($M_2 = (c_{1j})_{1\times t}$): *# Acquiring expectation of index intervals*

index_interval = 0

index_interval = random_choice ($[1, \cdots, t]$, probability=$M_2$) *# Pick one in random*

**Return** index_interval

*# Entire allocation in a single run*

**if __name__=='__main__':**

$\bar{n}$ = get_n_hat ($M_2 = (c_{1j})_{1\times t}$)

$x$ = [0,0,0,0,0,0,0,0,0,0,0,0,0,0,0,0,0,0,0,0]

**for all** j=[1, ⋯ ,20]

  **if** $b_{18j}$ is not null

    $x_j$=$b_{18j}$

**end for**

**for all** $k \in K$ **do**

  $G_k$= []

  i = k

  **while** i≥0

    i = i- positive_charge_index_interval ($M_2 = (c_{1j})_{1\times t}$)

    **if** i≥0

      $G_k$.add ( i )

  **end while**

  i = k

  **while** i < 20

    i = i + positive_charge_index_interval ($M_2 = (c_{1j})_{1\times t}$)

    **if** i < 20

      $G_k$.add ( i )

  **end while**

  **if** card ($G_k$) > 0

    allocate = $\frac{b_{18k}\bar{n}}{\mathrm{card}(G_k)}$

    **for all** j $\in G_k$ **do**

      **if** $LBDmj[j]$ *not in* $[D, E]$

        $x_j = x_j +$ allocate

      **if** $LBDmj[j]$ *in* $[D, E]$

        $x_j = x_j +$ 2×allocate

    **end for**

**end for**

**Supplementary Table 5.** Pseudocode of Algorithm 2.

**Algorithm 2** Relative-MIC value assignment of optimized LBD sequences

**Def** distance $(\beta(\{p_i^{\beta}\}_{i=1}^{2}),\{\alpha_j(\{p_i^{\alpha_j}\}_{i=1}^{2})\}_{j=1}^{k})$:

*# Require that* $\beta \epsilon K_2,\ \alpha_j \epsilon A$

$\{distance(\beta,\alpha_j)\}_{j=1}^{k} = 0$

**for all** $j = [1,\cdots,k]$ **do**

$\quad distance(\beta,\alpha_j) = \left(\sum_{i=1}^{2}(p_i^{\beta} - p_i^{\alpha_j})^2\right)^{\frac{1}{2}} \qquad \forall a \epsilon K_2$

**end for**

\# Require that $\mathrm{distance}(\beta,\alpha_1) \le \mathrm{distance}(\beta,\alpha_2) \le \cdots \le \mathrm{distance}(\beta,\alpha_k)$

**Return** $\{distance(\beta,\alpha_j)\}$

**Def** f $(\{relative\_MIC_a^i\},\ K_2)$:

$f(a,b) = 0$

$f(a,b) = \begin{cases} 1, & a > b \\ 0, & a \le b \end{cases}$

$\{f\left(relactive\_MIC_{\alpha_i}^{i}, relactive\_MIC_{\alpha_j}^{i}\right)\} \leftarrow \{relative\_MIC_a^i\} \quad i \ne j,\ \alpha_i, \alpha_j, \alpha \in K_2$

**Return** $\{f\left(relactive\_MIC_{\alpha_i}^{i}, relactive\_MIC_{\alpha_j}^{i}\right)\}$

**Def** get_kvalue ( $A = \{\alpha_j\}_{j=1}^{n}, \{actual\_MIC_{\alpha_j}\}_{j=1}^{n}$):

$\{\mathrm{competitivenessness\_index}_{\alpha}^{actual}\}_{\alpha \in A}, \{\mathrm{competitiveness\_index}_{\alpha}^{relative}\}_{\alpha \in A} = 0;\ \{Ind_i\}_{i=1}^{n-1} = 0$

**for all** $i = 1,\cdots,n-1$ **do**

$\quad$ **for all** $\alpha \in \mathrm{A}$ **do**

$\quad\quad \mathrm{competitiveness\_index}_{\alpha}^{actual} = \sum_{\alpha_i \in A} f\left(actual\_MIC_{\alpha_i}, actual\_MIC_{\alpha}\right)$

$\quad\quad \mathrm{competitiveness\_index}_{\alpha}^{relative} = \sum_{\alpha_i \in A} f\left(relactive\_MIC_{\alpha_i}^{i}, relactive\_MIC_{\alpha}^{i}\right)$

$\quad$ **end for**

$\quad Ind_i = \sum_{\alpha \in A}(\mathrm{competitiveness\_index}_{\alpha}^{actual} - \mathrm{competitiveness\_index}_{\alpha}^{relative})^2$

**end for**

$k \leftarrow \underset{i=\{1,2,\cdots,n-1\}}{i\_min} (Ind_i)$

**Return** $k$

**Def** relative_MIC $(\beta(\{p_i^{\beta}\}_{i=1}^{2}),\{\alpha_j(\{p_i^{\alpha_j}\}_{i=1}^{2})\}_{j=1}^{k},\ \{actual\_MIC_{\alpha}\}_{\alpha \in A})$:

*# Require that* $\beta \epsilon K_2,\ \alpha_j \epsilon A$

outputs, sum, k = 0, 0, 0

$k =$ get_kvalue ( $A = \{\alpha_i\}_{i=1}^{8}, \{actual\_MIC_{\alpha_i}\}_{i=1}^{8}$)

$relactive\_MIC_{\beta}^{k} = \sum_{i=1}^{k}\left(1 - \frac{distance(\beta,\alpha_i)}{sum}\right) \times actual\_MIC_{\alpha_i}$

**Return** $\{relactive\_MIC_{\beta}^{k}\}$

Note: $p_i$: $p_1$ is the net charge and $p_2$ is the hydrophobicity value.

**Supplementary Table 6.** Probe and primers used in this study.

| Primers | Sequence (5'-3') | Sequence information |
|---|---|---|
| WSSV32678-F | TGTTTTCTGTATGTAATGCGTGTAGGT | PCR |
| WSSV32753-R | CCCACTCCATGGCCTTCA | PCR |
| TaqMan probe WSSV32706 | CAAGTACCCAGGCCCAGTGTCATACGTT | qPCR |

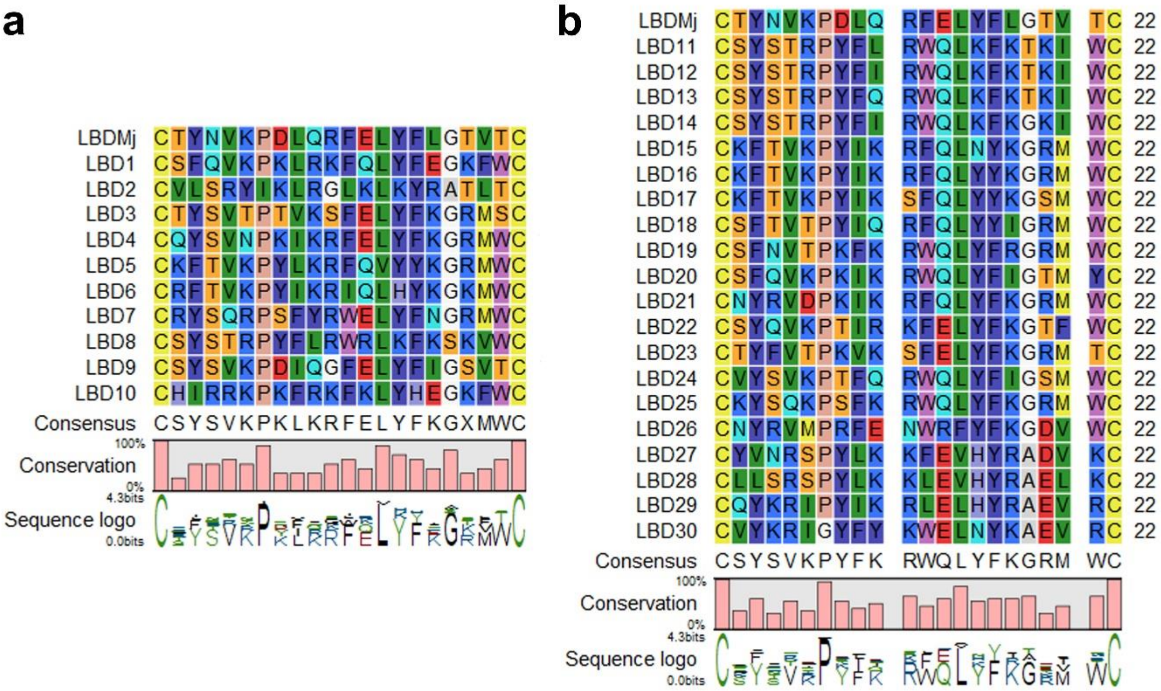


**Supplementary Figure 1.** Alignment of $LBD_{1\text{-}30}$ peptide sequences.

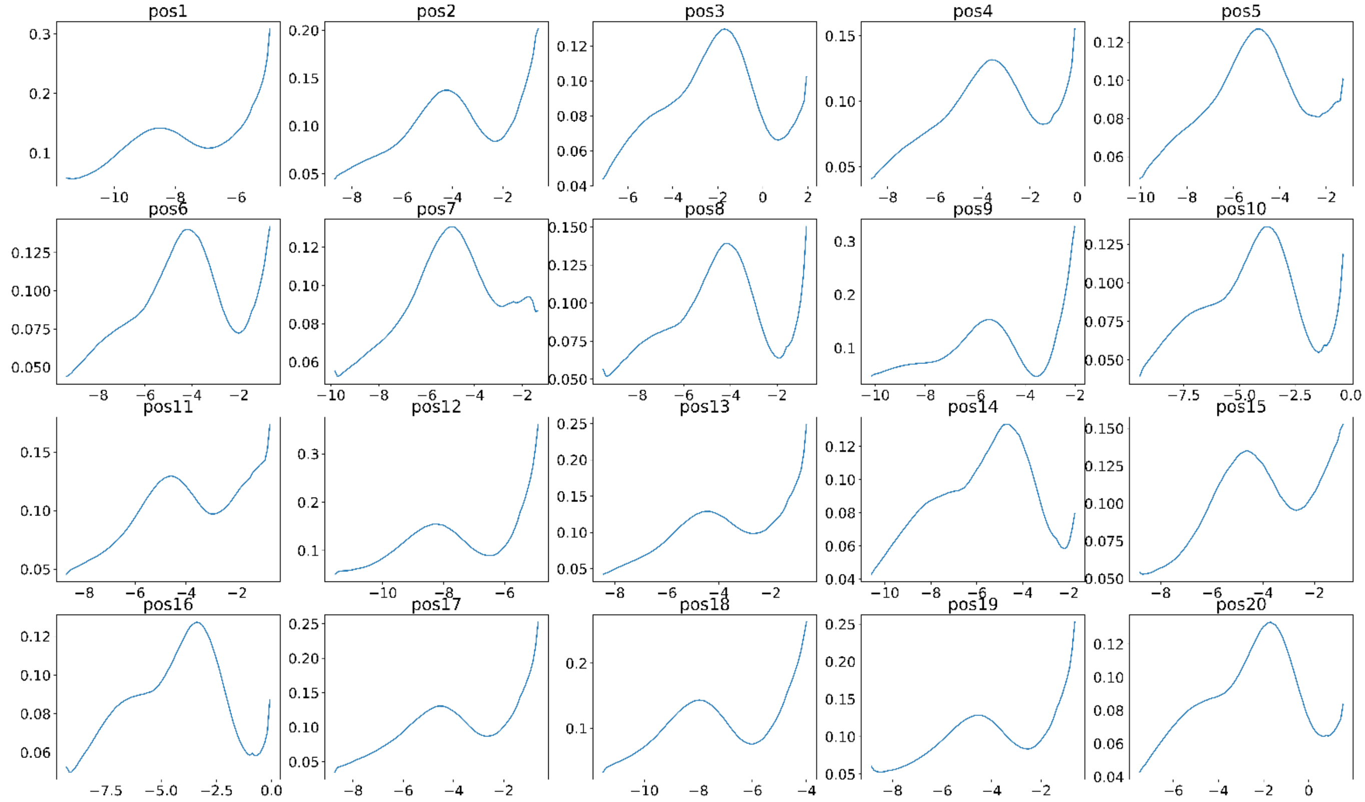


**Supplementary Figure 2.** Results of Monte Carlo simulation that indicates the probability distributions of the substitution coefficients of positively charged amino acids.

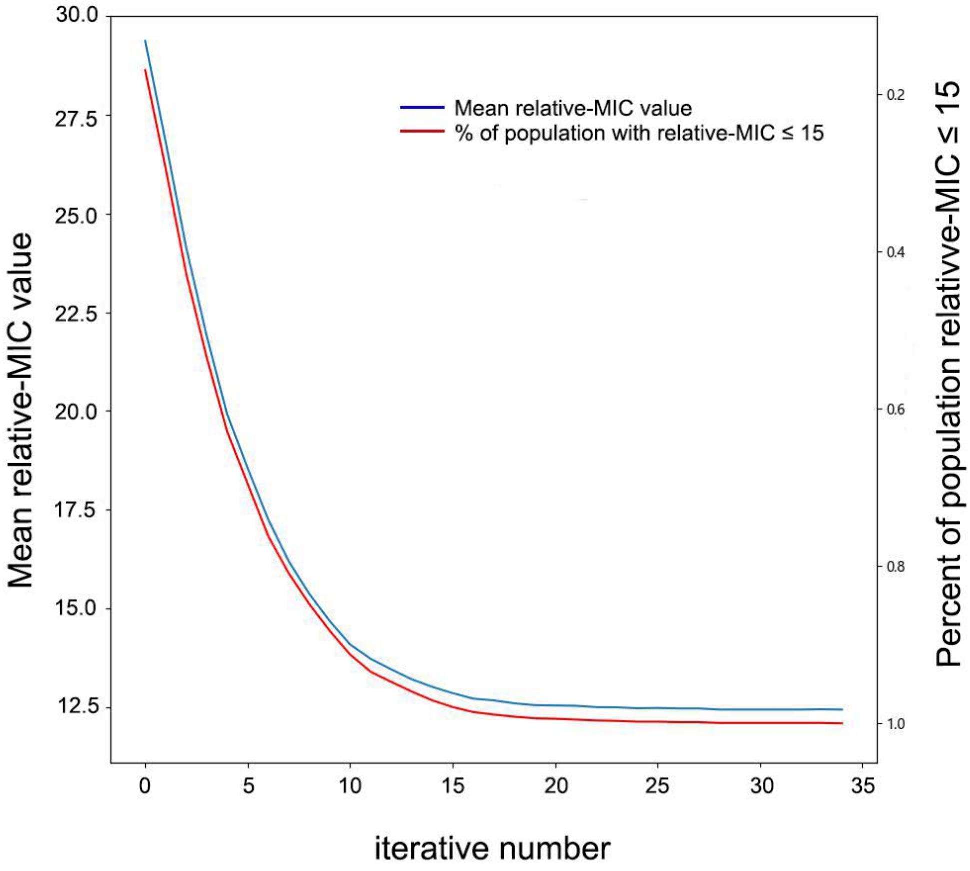


**Supplementary Figure 3.** Evaluation of the iteration number calculated by genetic algorithm in this work.

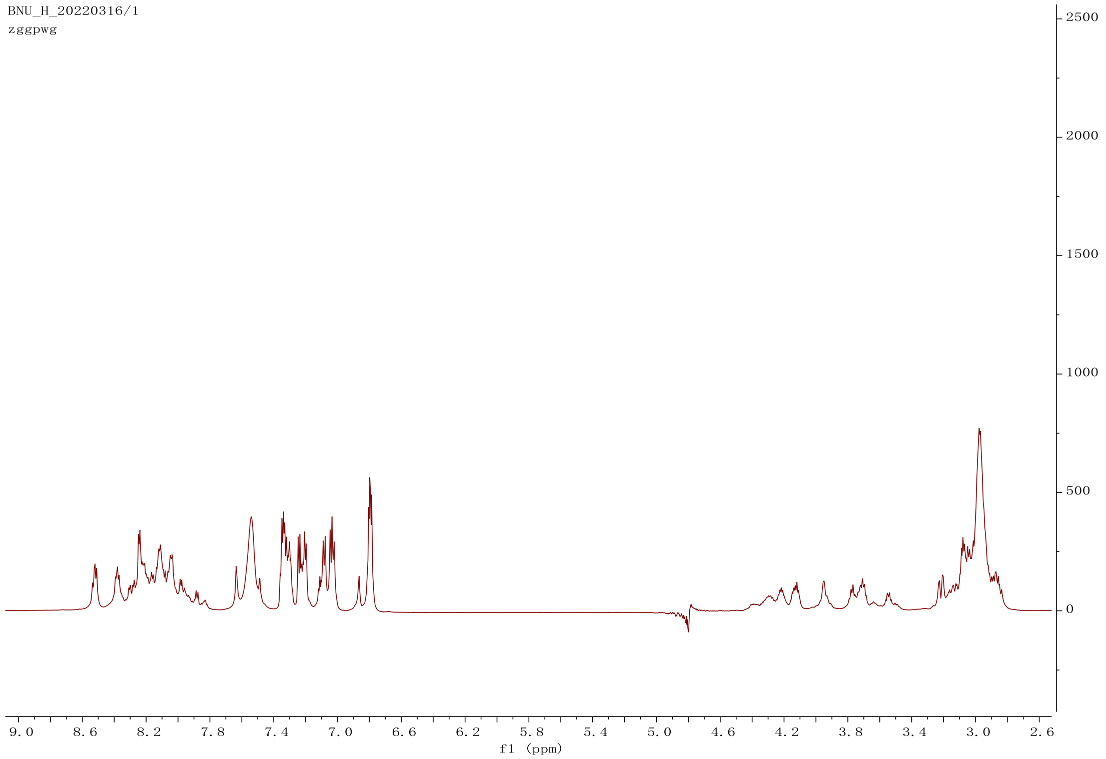


**Supplementary Figure 4.** $^{1}$H NMR spectrum for $LBD_B$. Condition: $H_2O$ with 10% $D_2O$, 700 MHz.

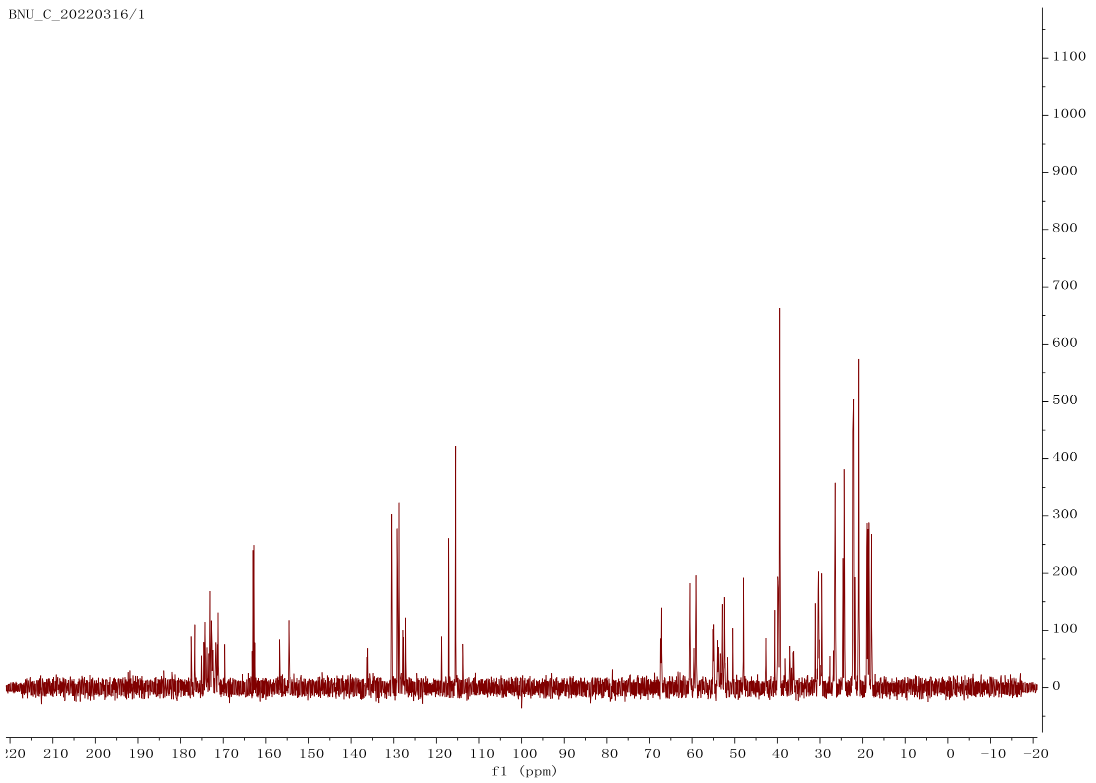


**Supplementary Figure 5.** $^{13}C$ NMR spectrum for $LBD_B$. Condition: $H_2O$ with 10% $D_2O$, 700 MHz.

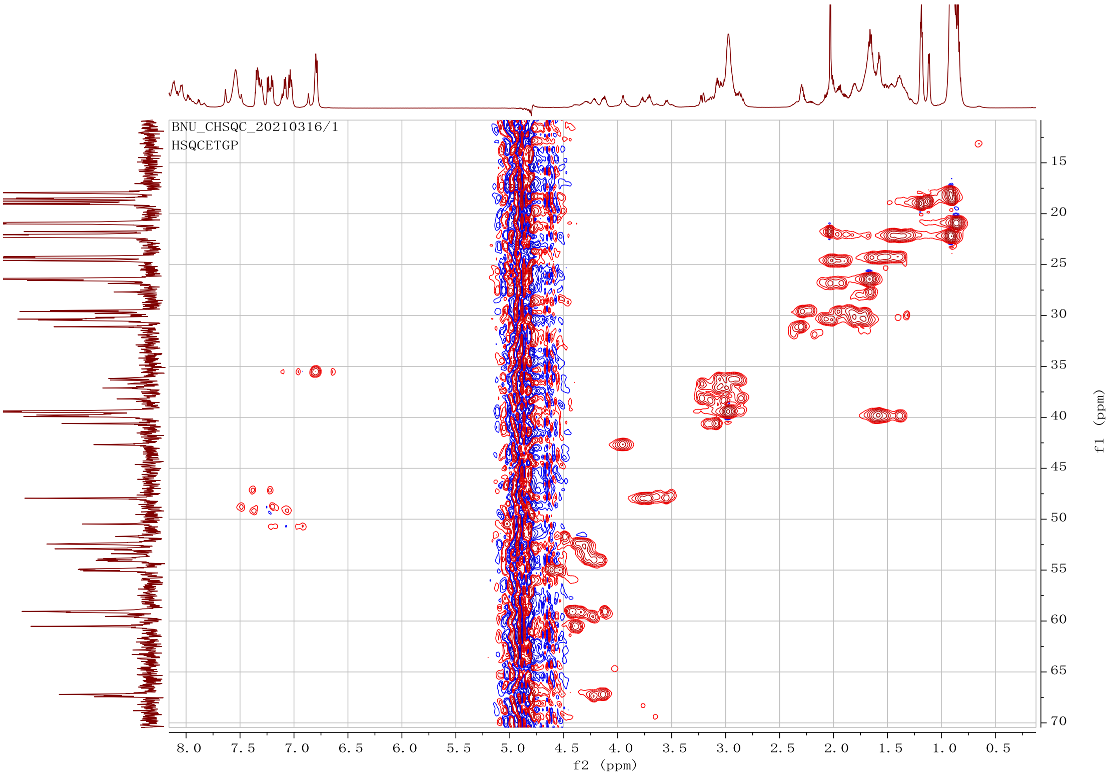


**Supplementary Figure 6.** HSQC NMR spectrum for $LBD_B$. Condition: $H_2O$ with 10% $D_2O$, 700 MHz.

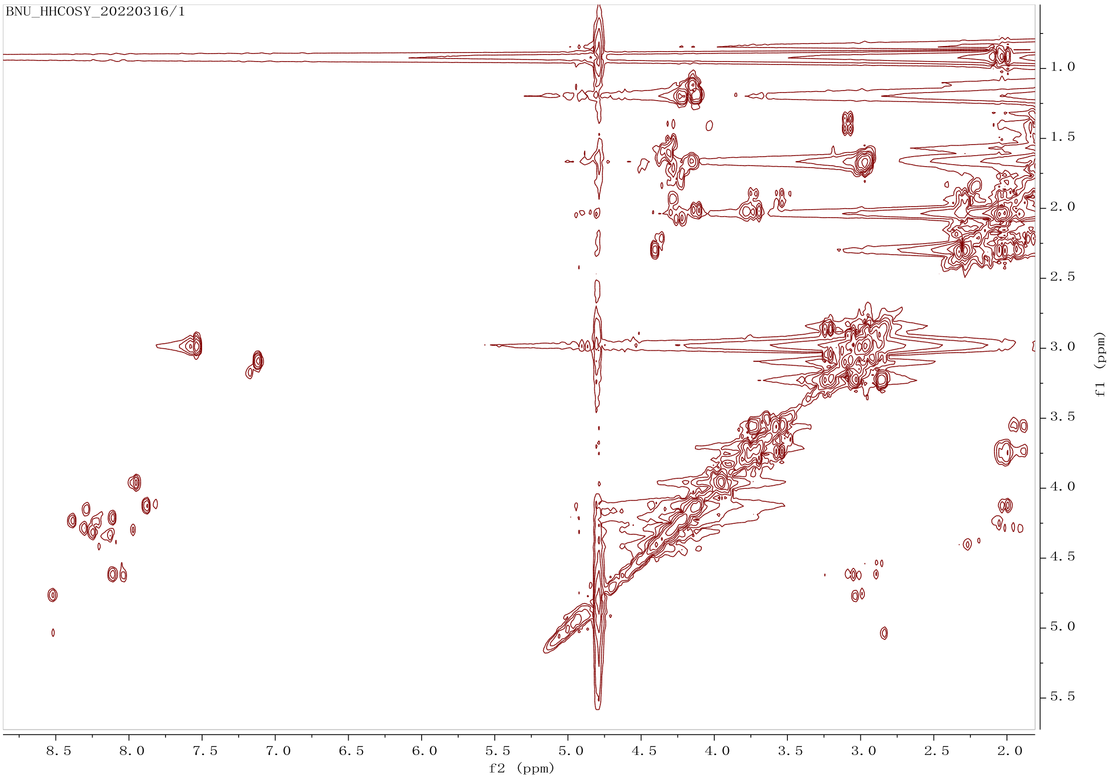


**Supplementary Figure 7.** $^1H$-$^1H$ COSY NMR spectrum for $LBD_B$. Condition: $H_2O$ with 10% $D_2O$, 700 MHz.

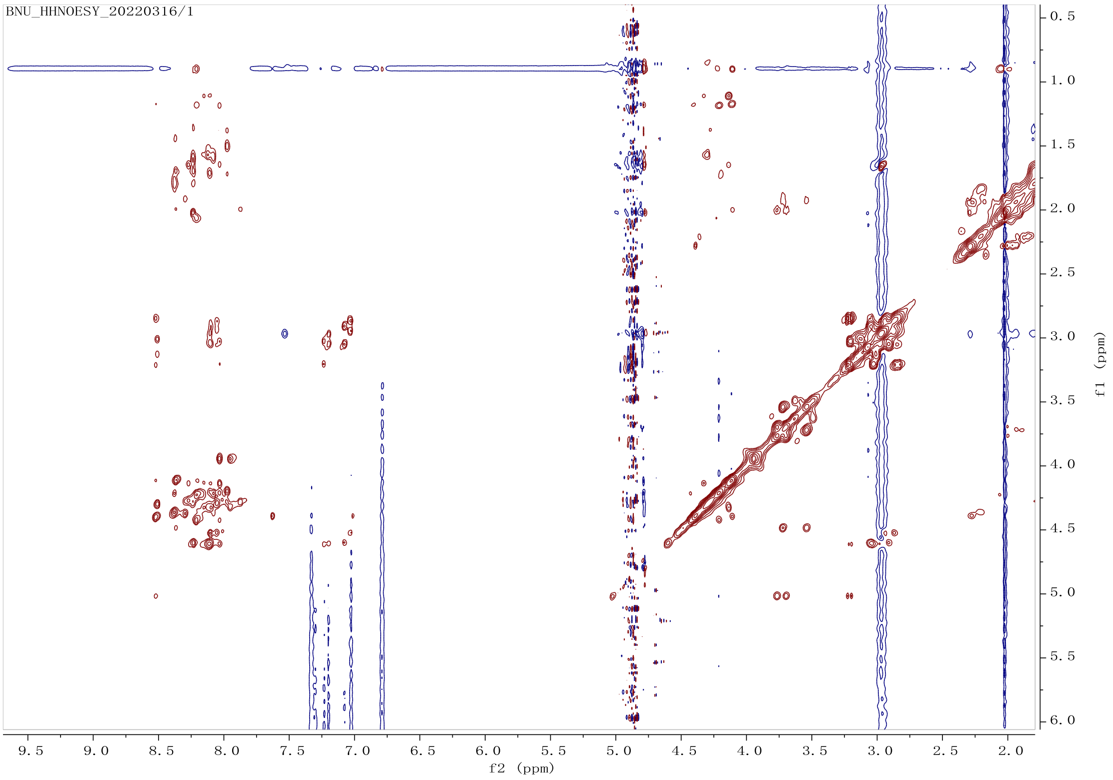


**Supplementary Figure 8.** NOESY NMR spectrum for $LBD_B$. Condition: $H_2O$ with 10% $D_2O$, 700 MHz.

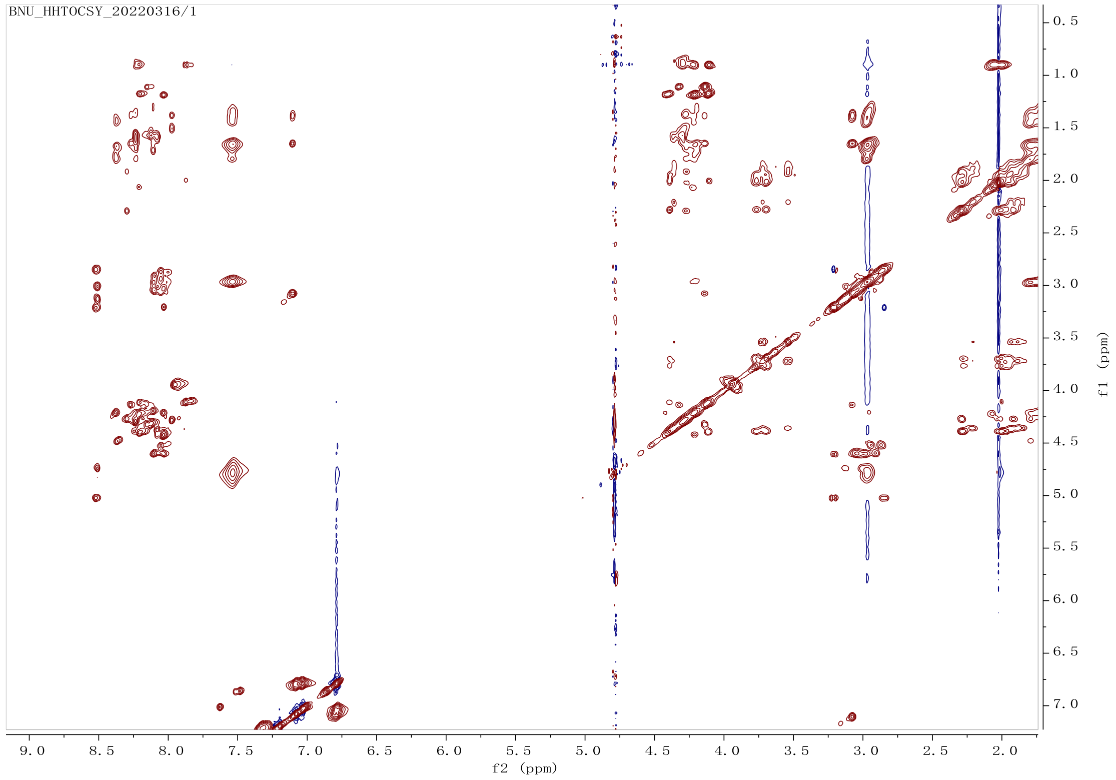


**Supplementary Figure 9.** TOCXY NMR spectrum for $LBD_B$. Condition: $H_2O$ with 10% $D_2O$, 700 MHz.
